# Perturbation response decomposition enables biologically aligned generalization to unseen perturbations and cellular contexts

**DOI:** 10.64898/2026.07.24.740459

**Authors:** Alexis Molina, Xinyi Zhang

**Affiliations:** AITHYRA, Research Institute for Biomedical Artificial Intelligence, Austrian Academy of Sciences, 1030 Vienna, Austria

## Abstract

Predicting single-cell responses to genetic perturbations could reveal the vast combinatorial space of perturbations and cellular contexts that is infeasible to measure experimentally, yet current deep learning models generalize poorly and often fail to outperform simple baselines. Here we demonstrate that generalizability in perturbation prediction requires identifying and representing distinct components of cellular response rather than on increasing model complexity alone. We introduce a decomposition framework that explicitly separates transcriptional responses into global, perturbation-specific, cell-line-specific, and perturbation-by-cell-line interaction components. Applied to four CRISPR-interference Perturb-seq screens on multiple cell lines, our framework reveals that these components have distinct structures and information requirements. The global response component is low-dimensional, reflects recurrent proliferation and stress response programs, and can be inferred from control gene expression. In contrast, the perturbation and cell-line specific components are high-dimensional and cannot be recovered from control expression. We therefore develop response-component-aligned models that map biological priors, such as gene coessentiality, onto the geometry of observed transcriptional responses. Critically, this alignment enables simple linear or multilayer perceptron (MLP) based models to outperform state-of-the-art architectures across multiple generalization settings, including unseen cell lines and combinations of unseen perturbations. Together, our framework for response decomposition and alignment provides a principled basis for evaluating and designing perturbation-prediction models, showing that generalization depends primarily on matching biological information to the response components rather than on model complexity alone.

## 1 Introduction

Large-scale perturbation assays, such as Perturb-seq, combine pooled CRISPR screens with single-cell RNA sequencing to map molecular interventions to transcriptome-wide phenotypes [1–5].These rich datasets enable insights into cellular functions, with implications for target prioritization, perturbation design, and mechanistic modeling. However, experimental screens can only sample a small fraction of the vast combinatorial space of cellular contexts and perturbations, making perturbation prediction a central challenge in computational biology. Perturbation prediction is a combinatorial generalization problem encompassing distinct, non-equivalent regimes, such as predicting an unseen perturbation, an unseen cell line, or an unobserved pairing of perturbation and cell line. These tasks impose different information requirements. A held-out perturbation in an observed cell line may be inferred from related perturbations or from transcriptional structures learned in that cell line, whereas cross-cell-line prediction requires estimating how a perturbation differentially affects an unseen cell line with different structures of gene programs than those represented in training. Understanding these basic axes of generalization is essential for model development and for extensions to combinatorial perturbations, chemical dose-responses, and full single-cell distributions.

Existing models use diverse representations and inductive biases to map cell states and interventions to transcriptional responses [6–11]. Recent advances use single-cell foundation models to learn generalized representations [9], model perturbation responses as cellular state transitions [10], or anchor predictions with external descriptors such as gene essentiality profiles [11]. These approaches not only differ in architecture, but their inputs and representations of perturbations and cellular contexts determine which components of a perturbation response they can predict in principle. However, increasing architectural complexity has not consistently improved predictive performance, with recent benchmarks suggesting that deep learning and single-cell foundation models underperform or merely match simple linear baselines [12]. It remains unclear whether this limitation reflects inadequate architectures, insufficient data, or a mismatch between a model’s inputs and the biological response it is expected to predict.

We reason that a major challenge is that standard aggregate metrics do not reveal which biological signals drive overall performance. The standard evaluation metrics can reward the recovery of systematic background variation or average responses rather than perturbation-specific effects [13]. A model may therefore obtain a high aggregate score by predicting a generic response across all perturbations, even if it fails to distinguish perturbation-specific effects or their dependence on cellular context. Model assessment should therefore ask not only which method attains the highest aggregate performance, but which cellular response components are predictable from the input data and recovered by the model.

Here we ask which components of a perturbation response are predictable from model inputs. We introduce a response decomposition framework that separates transcriptional responses into global, perturbation-specific, cell-line-specific, and perturbation-by-cell-line interaction components (Fig. 1a). This framework could guide the design of perturbation prediction models and decouple model capacity from data structure, revealing which biological signals are recovered underlying the overall performance. Across four CRISPRi Perturb-seq datasets [4, 5], we assess how control gene expression, prior biological knowledge, and observations of perturbed cell states constrain the predictability of each response component across different architecture and input choices. We identify a hierarchy of predictability. The global response component is low-dimensional, reflects recurrent proliferation and stress programs, and is represented within control expression variation. In contrast, perturbation-specific and cell-line specific components are higher dimensional and can only be predicted when biological priors that preserve functional relationships between genes are aligned with transcriptional response geometry. Whereas the perturbation-by-cell-line interaction component cannot be predicted without information from the target perturbation context. Guided by these findings, we demonstrate that simple response-aligned linear and MLP based models with gene coessentiality as priors [14] can predict the effects of unseen perturbations in unseen cell lines (Fig. 1b), outperforming more complex architectures. Extensions to combinatorial genetic and chemical perturbations further show that the most informative transferable representation depends on the perturbation modality. Together, response decomposition and alignment establish a comprehensive evaluation framework and guidance for model design, which explicitly reveal the predictable components in perturbation data and suggest that increasing model capacity alone does not necessarily lead to better generalization.

**Figure 1.**
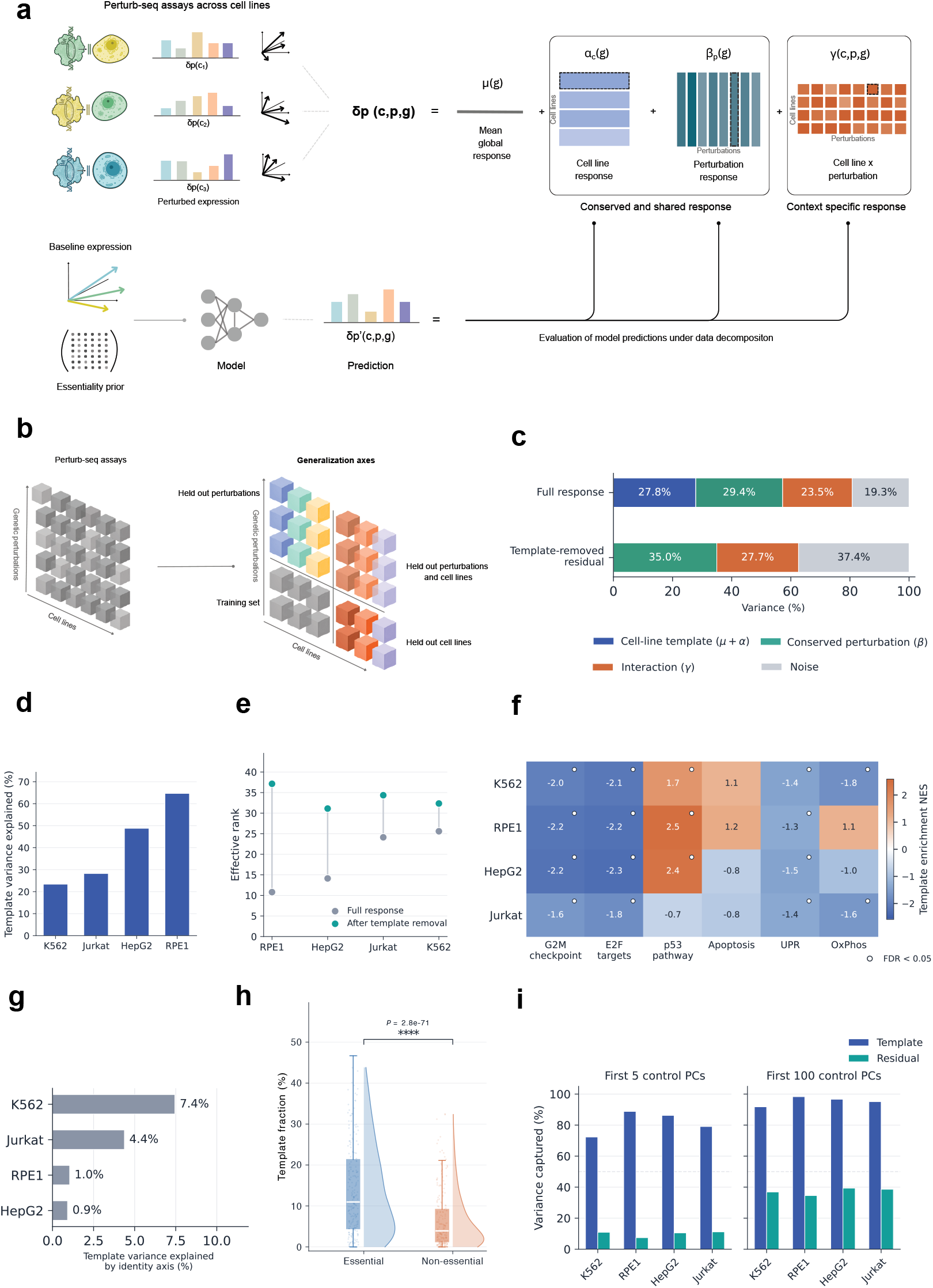
Perturbation responses decompose into shared response, conserved perturbation and context-specific components. a) Schematic of perturbation-response decomposition and model evaluation across cell lines. For each cell line *c*, perturbation *p*, and gene *g*, the pseudobulk response *δ*(*c, p, g*) was calculated as the difference between mean expression in perturbed cells and non-targeting controls. Responses were decomposed into a global mean response *µ*(*g*), a cell-line effect *α*_*c*_(*g*), a perturbation effect conserved across cell lines *β*_*p*_(*g*), and a cell-line-by-perturbation interaction *γ*(*c, p, g*). The sum *µ*(*g*) + *α*_*c*_(*g*) defines the mean response template for each cell line, whereas *β*_*p*_(*g*) and *γ*(*c, p, g*) capture conserved and context-specific perturbation effects, respectively. Models predict perturbation responses from baseline cell-state and perturbation information, and the same decomposition is applied to their predictions to determine which response components are recovered. b) Schematic of the different perturbation response prediction tasks. Within-cell-line prediction holds out perturbations within a given cell line. Cross-cell-line prediction additionally holds out the target cell line, while the most stringent setting holds out both the target cell line and the evaluated perturbations. c) Split half noise corrected decomposition of full and template removed responses across cell lines. Full responses contained a cell line template, a conserved perturbation component, a cell line by perturbation interaction and measurement noise. After template removal, residual responses retained both conserved perturbation signal and context specific interaction signal. d) Fraction of full perturbation response variance captured by the cell line specific template direction. Template contribution varied strongly across cell lines, with the largest template component in RPE1 and HepG2. e) Effective rank of full perturbation response matrices before and after template removal. Removing the shared template increased the effective dimensionality of the response space, indicating that the template captures a dominant low-dimensional component of the full response. f) Gene set enrichment analysis of cell line template directions. Colors show normalized enrichment scores for selected Hallmark signatures, and circles indicate FDR *<* 0.05. Cell line specific templates were enriched for recurring stress, proliferation and cell cycle related programs, with context dependent differences. g) Fraction of template variance explained by baseline cell identity axes derived from non targeting control cells. Baseline identity axes explained only a small fraction of the perturbation template, indicating that the template is not simply a restatement of cell line identity. h) Per-perturbation template fraction was computed by projecting responses onto the essential-gene template. Essential genes showed higher template contribution ((*p <* 0.0001) under a Mann-Whitney U test) than significant non-essential genes, indicating stronger and more consistent activation of the shared generic ^5^response program. i) Fraction of template and residual response variance captured by principal components of non targeting control expression. Control expression PCs captured much of the template direction but substantially less of the template removed residual, showing that baseline expression variation preferentially reflects shared response structure.

## 2 Results

### 2.1 Decomposition of perturbation response reveals predictability of response components

Predicting cellular responses to novel interventions requires generalizing from sparsely sampled combinations of cell line and perturbations to unseen perturbations, unseen cell lines, or completely new perturbation-cell line pairs. This remains a challenging problem that increasingly complex deep learning models do not consistently outperform simple linear baselines [12]. We reasoned that diagnosing modeling choices requires an explicit framing of the data structure that decouples the predictability of different response components from model capacity. We therefore develop a framework that decomposes perturbation responses into four orthogonal, noise-corrected components: a global mean response agnostic of cell line or perturbation, a cell-line-specific component, a perturbation-specific component conserved across cell lines, and a cell-line-by-perturbation interaction (Fig. 1a, Methods; Supplementary Fig. 1a,b; Supplementary Note 1). Applying this framework to each cell line in the Replogle-Nadig collections [4, 5], after noise correction, we found that reproducible variance was distributed across the cell-line-specific component (the cell-line template; 27.8%), the conserved perturbation-specific component (29.4%), and the cell-line-by-perturbation interaction (23.5%), with the remaining 19.3% attributable to measurement noise (Fig. 1c; Supplementary Fig. 1a,b). Even after removing the cell-line template, the interaction component dependent on both cell-line and perturbation accounts for up to 27.7% of the remaining variance (Fig. 1c; Supplementary Fig. 1a,b). Thus, perturbation responses consists of distinct layers. A substantial cell-line-agnostic component mixed with cell-line-specific and perturbation-specific components which remain context-dependent.

We next characterize how the cell-line templates align with the overall perturbation response geometry, which provides insights into why perturbation prediction models tend to mainly capture cell-line templates. The template’s prominence varied substantially across cell lines accounting for 23% to 65% of total response energy (Fig. 1d, Methods, Supplementary Note 2), reflecting how strongly each cell-line response space is dominated by a single direction aligned with the template (Supplementary Fig. 1c,d). After template removal, similar dimensionalities are observed across all four cell lines, indicating that apparent differences in response complexity largely reflect template magnitude rather than differences in core perturbation biology (Fig. 1e; Supplementary Fig. 1c). This explains why standard loss functions and full-response metrics often reward models that simply recover the cell-line template, even when perturbation-specific effects remain unresolved (Supplementary Note 1).

While the cell-line template represents a coordinated functional response, its magnitude and geometry depend both on cellular context and the function of the perturbed gene. Enrichment analysis shows the template is characterized by reproducible depletion of proliferation programs and enrichment of p53 and stress-associated pathways, though the specific pathways and magnitudes differed across cell lines (Fig. 1f; Supplementary Fig. 1f, Methods, Supplementary Note 2). Whereas cell line identity, derived from non-targeting control cells, explains only a small fraction of template variance (Fig. 1g). Although the cell-line template is conserved across perturbations targeting different genes, its magnitude is dependent on gene essentiality. Essential perturbations had approximately two-fold higher template contribution than significant non-essential perturbations (Fig. 1h, Methods), which persisted after response-magnitude matching (Supplementary Fig. 1g), even as the underlying biology remained highly concordant with depletion of E2F and MYC targets and enrichment of the p53 pathway (Supplementary Fig. 1h). This is consistent with the stronger template component observed in the non-transformed RPE1 compared with the cancer-derived K562, HepG2 and Jurkat cell lines, suggesting that essential-gene disruption engages a more coherent growth-arrest and stress-response program in a non-transformed cellular context (Fig. 1f). Finally, the perturbation-specific components not explained by the template used gene programs distinct from control-cell variation. The template was largely captured by the top principle components (PCs) of the control expression across all four lines, whereas the top 100 control PCs captured under 40% of residual variance, with the gap widening at with smaller number of PCs (Fig. 1i; Supplementary Fig. 1e; Supplementary Note 3). Our observations indicate that generalizing from unperturbed control cells alone is insufficient: the template’s dependence on cellular context and on perturbed-gene function, together with the perturbation-specific residual lying largely outside control-cell variation, shows that biological prior knowledge of both context and gene function is needed to identify perturbation-specific axes.

Together, these analyses separate three sources of perturbation response signal. First, perturbations induce a recurring cell-line template, shared across perturbations within each cell line, that is low-dimensional, biologically interpretable, and largely contained in control expression space. Second, perturbations induce template-independent residuals that are higher-dimensional and lie mostly outside the dominant control subspace, so that predicting the perturbation response using control cells alone requires biological priors to recover perturbation-specific axes. Third, across cell lines, much of the predictable residual signal reflects cell-line-by-perturbation interactions rather than perturbation effects conserved across cell lines. This decomposition therefore reframes perturbation prediction as a question of which response components are predictable given the experimental measurements, which we use throughout this study.

### 2.2 Response alignment makes gene coessentiality predictive of perturbation-specific effects

Generalization to held-out perturbations requires information about perturbation identity that baseline expression alone cannot provide. We therefore compared two representative approaches, a learned perturbation token such as in scGPT [9], and conditioning on biological prior knowledge such as using DepMap coessentiality profiles in MORPH [11]. scGPT mainly recovered the cell-line template whereas MORPH recovered perturbation-specific signal beyond the template (Fig. 2a), suggesting that external functional information about the perturbed gene is necessary for perturbation-specific prediction. However, the perturbation-specific response geometry, after template removal, is not directly encoded by similarity in DepMap profiles with close to zero correlations (Fig. 2b). Consistently, neither highly coessential gene pairs nor clusters in DepMap space showed substantial correlation with perturbation-specific response (Supplementary Fig. 2; Supplementary Note 4; Methods). This indicates that DepMap contains perturbation-relevant functional information, which is only predictive after response alignment. We demonstrate that this response alignment can be learned by simple models. Using matched five-fold perturbation-level cross-validation, Ridge and MLP models mapped projected DepMap profiles to pseudobulk responses, achieving comparable performance as the state-of-the-art model MORPH across all four cell lines (Fig. 2a; Methods; Supplementary Note 5). Thus, the recoverable signal arises primarily from combining an external functional prior with response alignment, rather than from conditional generative architecture.

**Figure 2.**
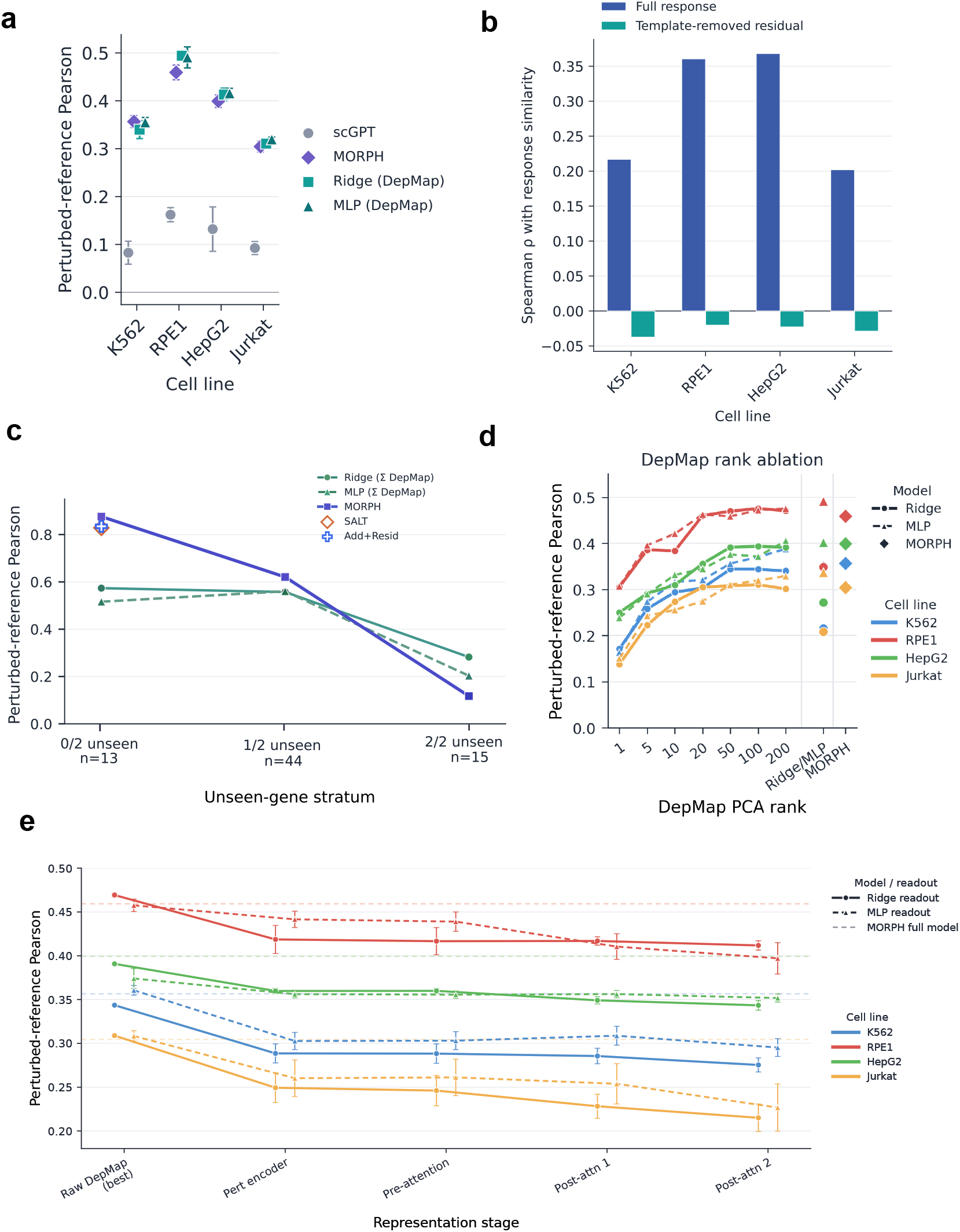
Response aligned transformations expose perturbation specific signal in coessentiality priors. a) Perturbation specific prediction performance within each cell line. Performance was evaluated using perturbed reference Pearson correlation in five fold perturbation level cross validation restricted to perturbations with available DepMap embeddings. Pairwise differences between models were assessed within each cell line using two-sided paired t-tests across the five matched cross-validation folds. scGPT performed significantly worse than MORPH, Ridge and MLP in all four cell lines (all (*p <* 0.001)). In RPE1, Ridge and MLP also significantly outperformed MORPH (*p* = 0.0031) and (*p* = 0.0106), respectively). In Jurkat, MLP significantly outperformed MORPH (*p* = 0.0231) and Ridge (*p* = 0.0475). No other pairwise differences were significant. b) Pairwise similarity between DepMap coessentiality profiles was compared with pairwise similarity between perturbation responses. Raw DepMap similarity was positively associated with full response similarity but not with template removed residual similarity, indicating that raw coessentiality neighborhoods do not define residual response neighborhoods. c) Norman double-gene perturbation benchmark stratified by single-gene exposure. Test double perturbations were grouped by the number of constituent genes absent from the single-gene training set. Ridge and MLP baselines were trained on double-perturbation training examples using pairwise DepMap dependency features. SALT and Add+Resid are shown only for the 0/2 unseen setting, where both constituent single-gene responses are available. MORPH performed best when single-gene effects were observed, but dropped sharply in the 2/2 unseen setting, whereas raw DepMap baselines retained predictive signal d) DepMap PCA rank ablation. Ridge and MLP readouts were trained using increasing numbers of DepMap principal components and evaluated by perturbed-reference Pearson. Performance rose rapidly at low rank and saturated around 20–50 PCs across cell lines, indicating that response-aligned signal is concentrated in a compact coessentiality basis. e) MORPH representation-stage analysis. Predictive content was compared across the raw DepMap profile, perturbation encoder representation, pre-attention representation and post-attention representations. The raw DepMap profile was the most predictive representation in all four cell lines. The MORPH perturbation encoder reduced performance, the pre-attention representation did not add information beyond the perturbation encoder, and downstream attention layers did not reco_6_ver the lost signal. This shows that MORPH’s predictive signal is carried primarily by the DepMap descriptor rather than by progressive response alignment through the conditional architecture.

For combinatorial perturbation prediction, better response alignment with simple models improves generalization compared to state-of-the-art models. Using the Norman dataset [15], our benchmark showed that MORPH matched additive baselines [16] when both constituent genes had been seen as single perturbations during training, but this advantage diminished when one gene was unseen and reversed when both were unseen, with MORPH underperforming simpler models (Fig. 2c). In this setting of combination of unseen perturbations, Ridge and MLP models trained on double perturbations retained predictive signal from pairwise DepMap features, indicating that MORPH relies on measured single-gene effects, whereas response-aligned coessentiality maps generalize more effectively to novel gene pairs.

To understand why simple models match or outperform a more complex deep learning model, we inspected how each model converts the coessentiality prior into a predictive perturbation representation. Rank ablation over the DepMap PCA projection showed that the learnable signal is low-rank, with Ridge and MLP performance saturating between 20-50 principal components across all four cell lines (Fig. 2d; Methods, Supplementary Note 6-7). Thus, perturbation-specific prediction does not require the full dependency profile, but a low-dimensional set of coessentiality programs aligned with template-removed transcriptional responses. We then asked whether MORPH learned a more informative transformation of this prior. Unlike Ridge and MLP, which map DepMap directly into response space, MORPH first compresses the dependency profile before integrating it with control expression through downstream attention modules. Applying the same response-aligned readout to intermediate MORPH representations showed that the raw DepMap profile was the most predictive representation in every cell line. MORPH’s perturbation encoder reduced recoverable signal, and subsequent integration and attention layers did not improve alignment with the response geometry (Fig. 2e; Supplementary Note 8; Methods). Together, these analyses indicate that the predictive DepMap prior is low dimensional and best recovered through direct response alignment, whereas increasing model complexity can result in worse alignment and generalization performance rather than revealing additional perturbation-specific structure. Thus, the central modeling problem is not architectural complexity alone, but learning an operator that converts dependency variation into the template-removed response component being predicted.

### 2.3 Mapping perturbation-aligned axes across cell lines enables generalization to unseen cell lines

Out-of-context prediction in a held-out cell line requires learning perturbation effects that transfer across contexts while decoding them into a target response geometry that has not been directly measured. While existing models claim the ability to generalize to unseen cell lines [10, 11], our decomposition framework shows that these models cannot predict perturbation-specific components in held-out cell lines beyond the cell-line template (Fig. 3a; Supplementary Fig. 4b; Methods, Supplementary Note 9). This lack of generalizability of existing models by using control expression of the target cell line is expected, given our decomposition results that the top PCs of the control cells do not capture variance beyond cell-line template (Fig. 1g). Without a direct measurement of perturbation in the target cell line, we demonstrate that out-of-context prediction is achievable by learning a mapping between the perturbation-aligned bases of the source and target response space through prior knowledge. We trained Ridge and MLP models to map perturbation measurements from three source cell lines, using projected DepMap profiles as input descriptors, and evaluated them in the held-out fourth cell line. Both models were able to predict template-removed signals in every held-out cell line, with MLP outperforming Ridge but the same ordering of cell lines in terms of prediction performance (Fig. 3a, Supplementary Note 10-11). Incorporation of more source cell lines improved generalization performance for both models and in all held-out cell lines (Fig. 3a). Importantly, our framework using either Ridge or MLP enables prediction of cell line template-removed signal even when both the target cell line and the perturbed gene were held-out from training (Fig. 3b, Supplementary Note 11).

**Figure 3.**
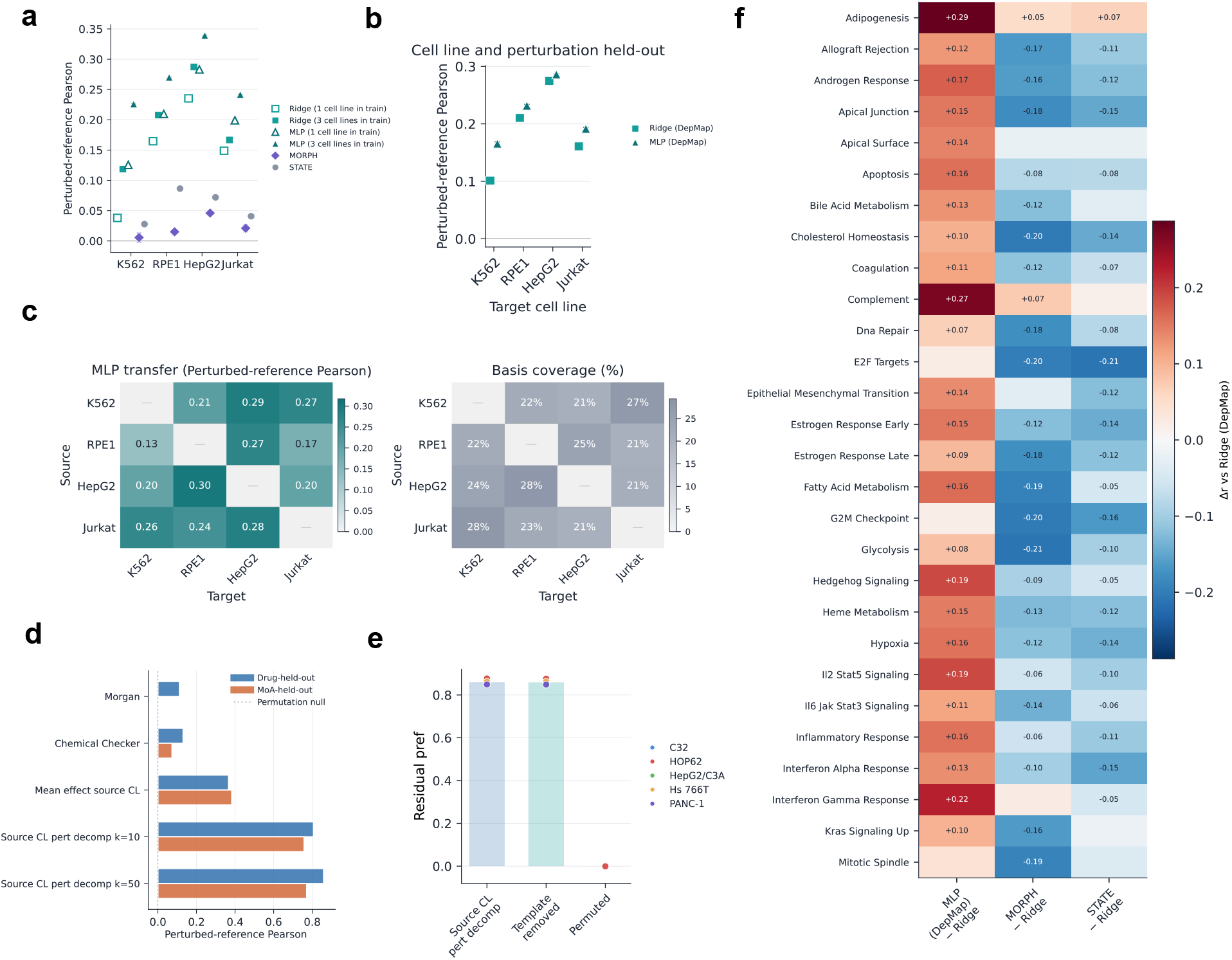
Source responses and coessentiality priors reveal transferable perturbation signal across contexts. a) Cross-cell-line transfer for genetic perturbations. Models were trained using either one source cell line or three pooled source cell lines and evaluated in a held-out target cell line. DepMap-based Ridge and MLP models recovered positive template-removed signal across all targets, with consistently higher performance when three source cell lines were used. For MLP, the improvement from one to three source cell lines was significant in every target cell line (two-sided Welch’s t-test across five cross-validation folds, (*p <* 10^−10^) for all comparisons). By contrast, MORPH and STATE recovered little template-removed signal in the source-only setting. b) Double-unseen transfer, where both the target cell line and evaluated perturbation genes were held out during training. DepMap-based Ridge and MLP retained positive perturbed-reference Pearson across all target cell lines, indicating that coessentiality profiles encode perturbation information that generalizes beyond memorized source perturbations. c) Directed one-source to one-target transfer and source-response basis coverage. Left, MLP perturbed-reference Pearson for each directed transfer pair, where MLP was trained using the source cell line to predict perturbation effects in the target cell line. Right, fraction of target template-removed residual variance spanned by the source response basis at rank 20. Source bases covered only a limited fraction of target residual variance and did not track transfer performance, indicating that DepMap-based models do not simply reuse source transcriptional response geometry. d) Chemical perturbation transfer in Tahoe-100M. Static compound priors, including Morgan fingerprints and Chemical Checker signatures, weakly predicted template-removed chemical responses, whereas observed source-cell-line responses were substantially more informative. Low-rank Ridge models trained on source-response programs strongly outperformed static compound priors in both held-out-compound and leave-one-MoA-out evaluations. e) Controls for source-response transfer in Tahoe-100M. Source-response Ridge retained high performance after template removal and collapsed under compound-label permutation, showing that transfer was not driven by a generic template or leakage but by chemical-specific response coordinates observed in source cell lines. Points show target-cell-line results. f) Pathway-level recovery in cross-cell genetic perturbation prediction. True and predicted template-removed responses were projected onto Hallmark pathway vectors, and pathway activity prediction was compared with Ridge on DepMap. MLP improved recovery of selected stress, immune and signaling programs, whereas MORPH and STATE generally lost pathway-level signal relative to the raw DepMap Ridge baseline. This indicates that cross-context prediction is best explained by recovery of response-aligned perturbation coordinates rather than by model-specific pathway targeting.

The difference between perturbation-aligned bases of the source and target response spaces indicates difference in core perturbation-induced gene programs in different cell lines. We define perturbation-aligned bases as the top singular vectors of a cell line’s template-removed perturbation response matrix. By projecting the target cell line’s perturbation response matrix to the bases of the source cell line, we quantified that the source cell line’s bases only capture 20% to 30% of target variance at rank 20 after template removal (Fig. 3c; Supplementary Fig. 4c–e, Supplementary Note 12-14). Additionally, the extent of alignment in perturbation bases does not correlate with model performance (Fig. 3c). Thus, coessentiality provides an external functional coordinate system that maps perturbation-aligned bases of the source and target cell lines’ response spaces, despite limited overlap between their expression programs.

Having established that generalization to unseen genetic perturbations and unseen cell lines can be achieved through biological prior, we explored out-of-context generalization for chemical perturbations where no analogous prior is available. We tested generalization to unseen chemical perturbations, with either zero-shot learning through chemical descriptors encoding molecular identity or few-shot learning through the target perturbation response measured in a source cell line encoding an empirical, one-shot prior. In both settings, the compound was excluded from model training across all cell lines. Using Tahoe-100M [17], we compared Morgan fingerprints [18] and Chemical Checker signatures [19] with drug responses observed in other cell lines (Methods). Static descriptors poorly predicted held-out compound effects, whereas one-shot transfer based on source cell lines had substantially better performance, with a low-rank Ridge regression over source-response programs performed best (mean r=0.86, compared with r=0.11 for Morgan fingerprints; Fig. 3d). This better performance of few-shot transfer persisted when entire mechanisms of action were held out. Although compound responses also contained a cell-line template, one-shot transfer performance remained strong after template removal and collapsed under compound-label permutation, showing that prediction depended on matched chemical-specific response coordinates rather than shared response structure alone (Fig. 3e). Thus, when an external functional prior is unavailable, perturbation responses measured in alternate cellular contexts can provide an empirical prior for out-of-context prediction.

Finally, we assessed generalization performance of each model at the pathway-level for any variability in performance associated with gene functions that would not be captured by an aggregate score. We projected true and predicted template-removed responses onto Hallmark pathway vectors and measured concordance across perturbations (Methods). While both using DepMap as the biological prior, the MLP model shows an overall improvement compared to the Ridge regression and the improvement is the highest for signaling, inflammatory and stress-associated programs (Fig. 3f). Whereas MORPH [11] and STATE [10], which do not use prior, performed worse across all pathways compared to the DepMap Ridge regression baseline (Fig. 3f). Thus, pathway-level analysis showed that nonlinear mapping of perturbation response axes can improve generalization to unseen cell lines, but increased architectural complexity alone does not improve generalizability of any pathways.

In summary, out-of-context generalization depends on both the source of transferable perturbation information and the model used to learn the mapping. Coessentiality provides an external functional prior for genes, whereas measured source-cell-line responses provide an empirical prior for compounds. In both settings, successful prediction depends on transforming transferable perturbation information into response-aligned coordinates, while increased architectural complexity alone does not expose additional cross-context or pathway-level signal and could result in worse performance.

### 2.4 Our decomposition framework reflects the predictability of the response components

Performance metrics used to evaluate perturbation prediction models usually rely on aggregate scores, which do not reflect the performance on predicting each biological component [13]. Our framework demonstrates that models can differ substantially in recovering the global, perturbation-specific, cell-line-specific (template), and cell-line-by-perturbation interaction components, even at similar aggregate performance. We therefore applied the response decomposition to model outputs at evaluation time. For generalization to unseen perturbation, we separated predictions into template-aligned and template-removed components. For generalization to unseen cell lines, we measured the alignment of model predictions with the perturbation-specific component conserved across cell lines *β*_*p*_(*g*), and the cell-line-by-perturbation interaction component, *γ*(*c, p, g*) (Fig. 4; Supplementary Fig. 5; Supplementary Note 16; Methods).

**Figure 4.**
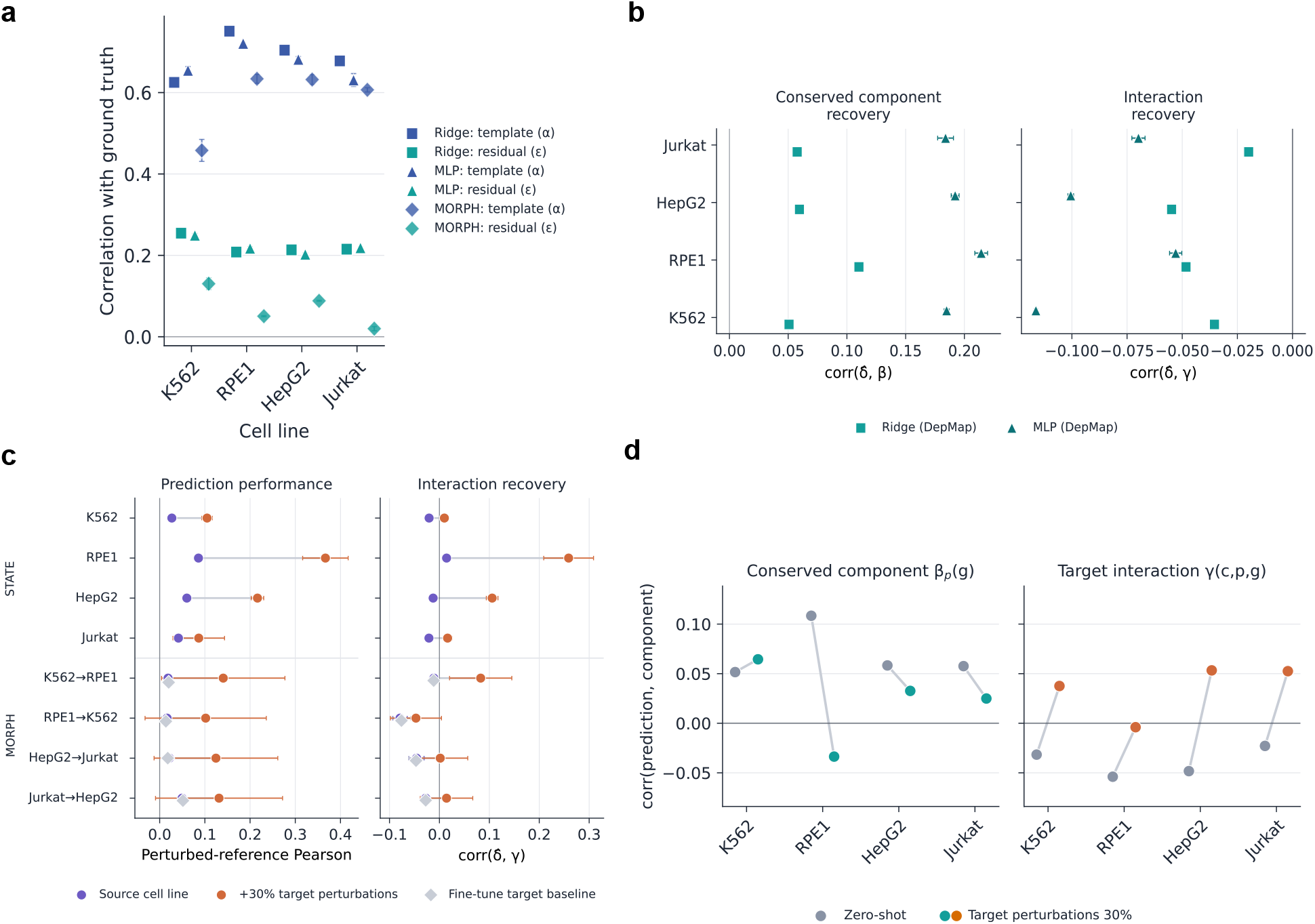
Prediction decomposition exposes transferable signal and target specific interaction limits. a) Within-cell-line prediction attribution. True and predicted responses were decomposed into template-aligned coefficients and template-removed residuals. Ridge, MLP and MORPH recovered template coefficient variation more strongly than residual variation, indicating that positive prediction performance contains substantial shared response structure together with weaker perturbation-specific residual recovery. b) Cross-cell-line zero-shot prediction attribution for DepMap-based Ridge and MLP models. Predictions were compared with the conserved perturbation component (*β*_*p*_(*g*)) and the target interaction component (*γ*(*c, p, g*)) from the balanced cell-line-by-perturbation decomposition. Models aligned positively with the conserved component but not with the target interaction, showing that source responses and DepMap recover transferable perturbation signal but not the target-specific interaction. c) Effect of target perturbation calibration on STATE and MORPH. Source-only transfer produced weak perturbed-reference performance and little or negative interaction recovery. Adding 30% target perturbations improved performance and shifted predictions toward positive interaction recovery, whereas MORPH fine-tuning on target controls alone did not recover the missing interaction. d) Target calibration baseline decomposition. Training with 30% target perturbations increased alignment with the target interaction (*γ*(*c, p, g*)), while recovery of the conserved component (*β*_*p*_(*g*)) changed little or decreased. These analyses show that target perturbation measurements improve prediction primarily by revealing the cell-line-by-perturbation interaction rather than by increasing recovery of the conserved perturbation component.

**Figure 5.**
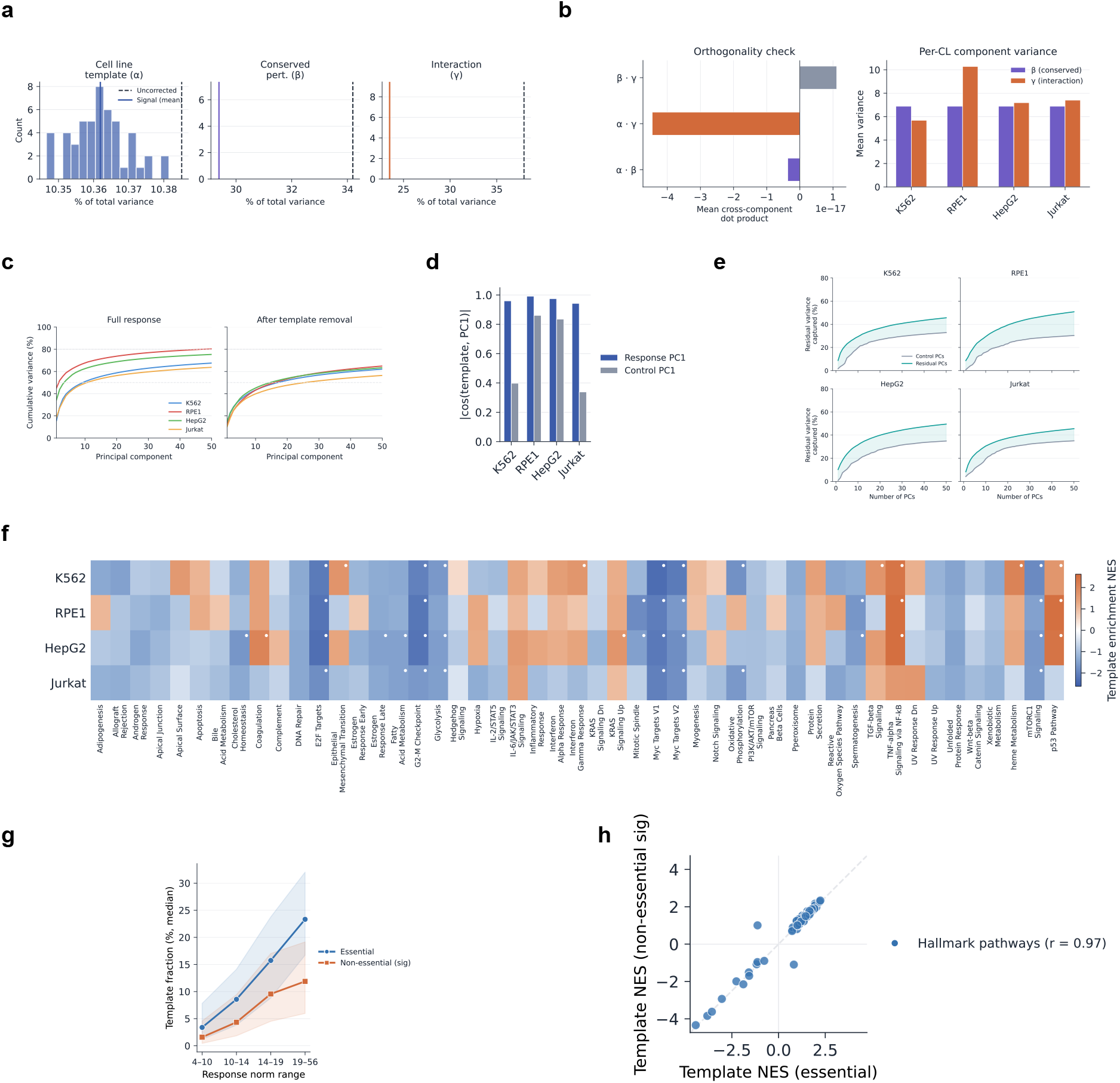
Template and residual geometry of perturbation responses. a) Split-half noise correction for the full-response decomposition. Histograms show uncorrected component variance estimates across resampling splits. Vertical solid lines show the mean reproducible signal estimate from cross-half dot products, and dashed lines show the corresponding uncorrected estimates. b) Additive decomposition diagnostics. The left panel shows mean cross-component dot products, confirming near orthogonality of the cell-line, conserved perturbation and interaction components in the balanced response tensor. The right panel shows per-cell-line component variances, indicating that both conserved perturbation effects and cell-line-by-perturbation interactions contribute to reproducible response structure. c) Cumulative variance explained by principal components of the full pseudobulk perturbation response matrix and the template-removed residual response matrix for each cell line. Full responses showed stronger low-dimensional structure in template-dominated cell lines, whereas residual responses had more similar spectra across cell lines after removal of the shared template direction. d) Absolute cosine similarity between the cell-line template and the first principal component of either the perturbation response matrix or the non-targeting control expression matrix. The template aligned strongly with the leading perturbation-response axis but was not equivalent to the leading control-expression axis. e) Fraction of residual response variance captured by principal components fitted either to non-targeting control cells or to template-removed perturbation residuals. Control PCs captured only a limited fraction of residual variance compared with residual PCs, indicating that perturbation-specific residual structure is only partly visible in baseline expression variation. f) Hallmark gene-set enrichment analysis of cell-line-specific template directions. Templates showed recurrent enrichment and depletion of stress, proliferation and cell-cycle-associated programs, with context-dependent differences across K562, RPE1, HepG2 and Jurkat. g) Per-perturbation template fraction was computed by projecting each response onto the essential-gene template and measuring the fraction of response energy captured by that direction. Perturbations were stratified by response-magnitude quartiles and compared between essential genes and significant non-essential genes. Essential perturbations showed higher template contribution across matched response-strength bins, indicating that their stronger template loading is not explained solely by larger response magnitude. h) Hallmark gene-set enrichment was computed separately for the essential-gene template and the significant non-essential-gene template. Each point represents one Hallmark pathway, with normalized enrichment scores compared between the two templates. The strong concordance shows that essential and significant non-essential perturbations activate the same generic response program, rather than distinct template biology. Essential perturbations therefore differ mainly in template strength and consistency, not in template identity.

**Figure 6.**
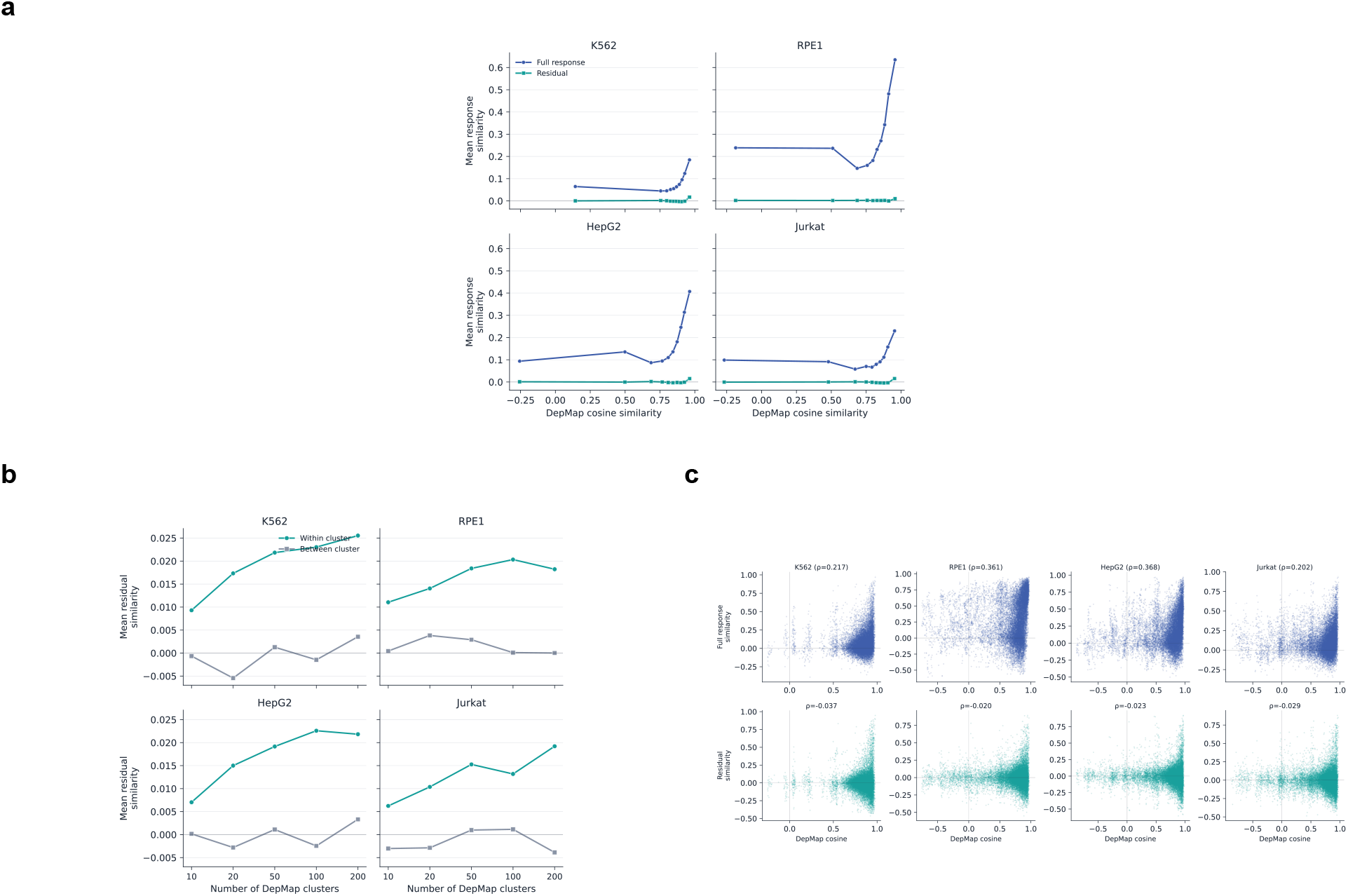
Raw coessentiality similarity does not define template removed response geometry. a) Mean response similarity as a function of DepMap cosine similarity decile for each cell line. Raw DepMap similarity was associated with full response similarity. This association was largely lost after template removal. b) Residual response coherence within DepMap clusters. Genes were clustered in DepMap coessentiality space using different numbers of clusters. Mean template removed response similarity was compared within clusters and between clusters. Within cluster residual similarity was only modestly higher than between cluster similarity. This indicates that DepMap neighborhoods do not directly define residual response neighborhoods. c) Pairwise relationship between DepMap cosine similarity and perturbation response similarity. Each point represents a pair of perturbation genes. Full response similarity showed a positive association with DepMap similarity. Template removed residual similarity showed weak or near zero association across K562 RPE1 HepG2 and Jurkat.

**Figure 7.**
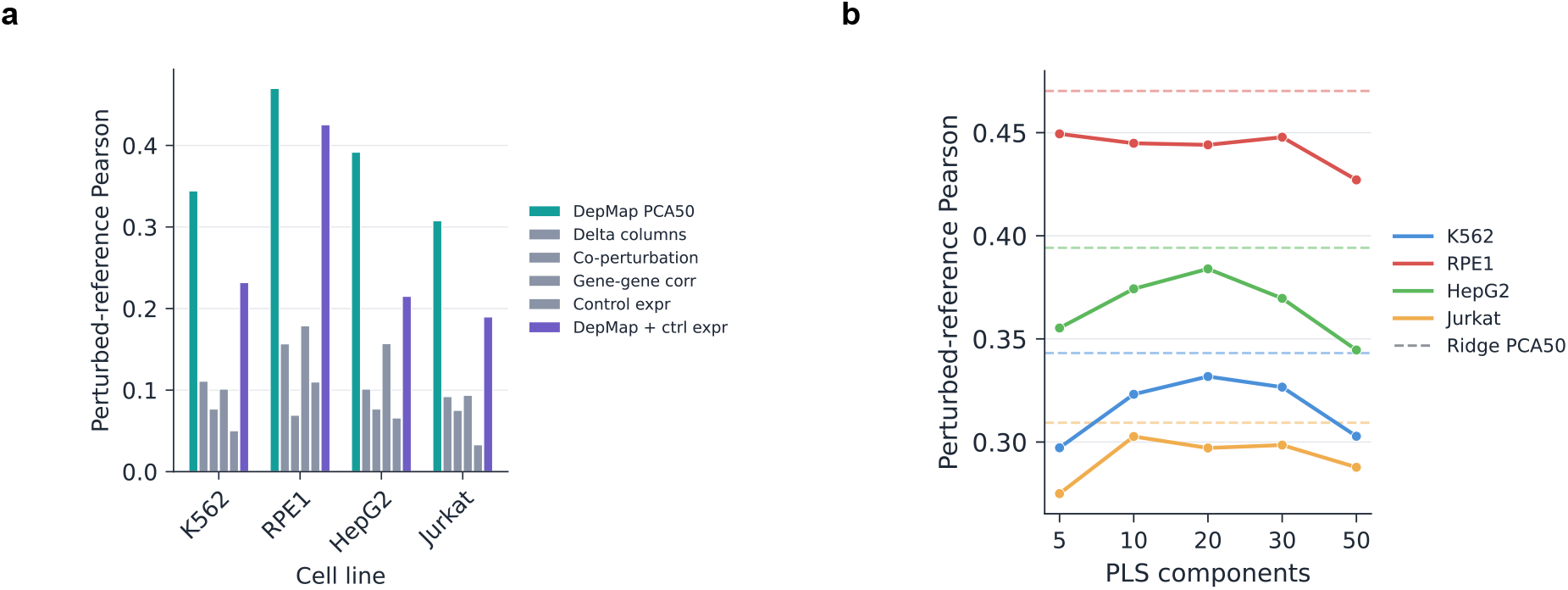
Descriptor ablations and response aligned retrieval controls. a) Perturbation prediction from alternative gene descriptors. Bars show perturbed reference Pearson correlation for each cell line. DepMap PCA50 outperformed expression derived descriptors including delta columns co perturbation profiles gene gene expression correlations and control expression statistics. Adding control expression statistics to DepMap did not consistently improve performance. This indicates that the predictive signal is primarily carried by the external coessentiality prior rather than by descriptors derived from the expression matrix. b) Partial least squares response aligned retrieval from DepMap profiles. Curves show perturbed reference Pearson correlation as a function of the number of PLS components. Dashed horizontal lines show the corresponding Ridge PCA50 baseline for each cell line. PLS based retrieval recovered positive perturbation specific signal but generally remained below the supervised Ridge map. This supports the conclusion that DepMap contains transferable perturbation information that becomes useful after alignment to response geometry.

**Figure 8.**
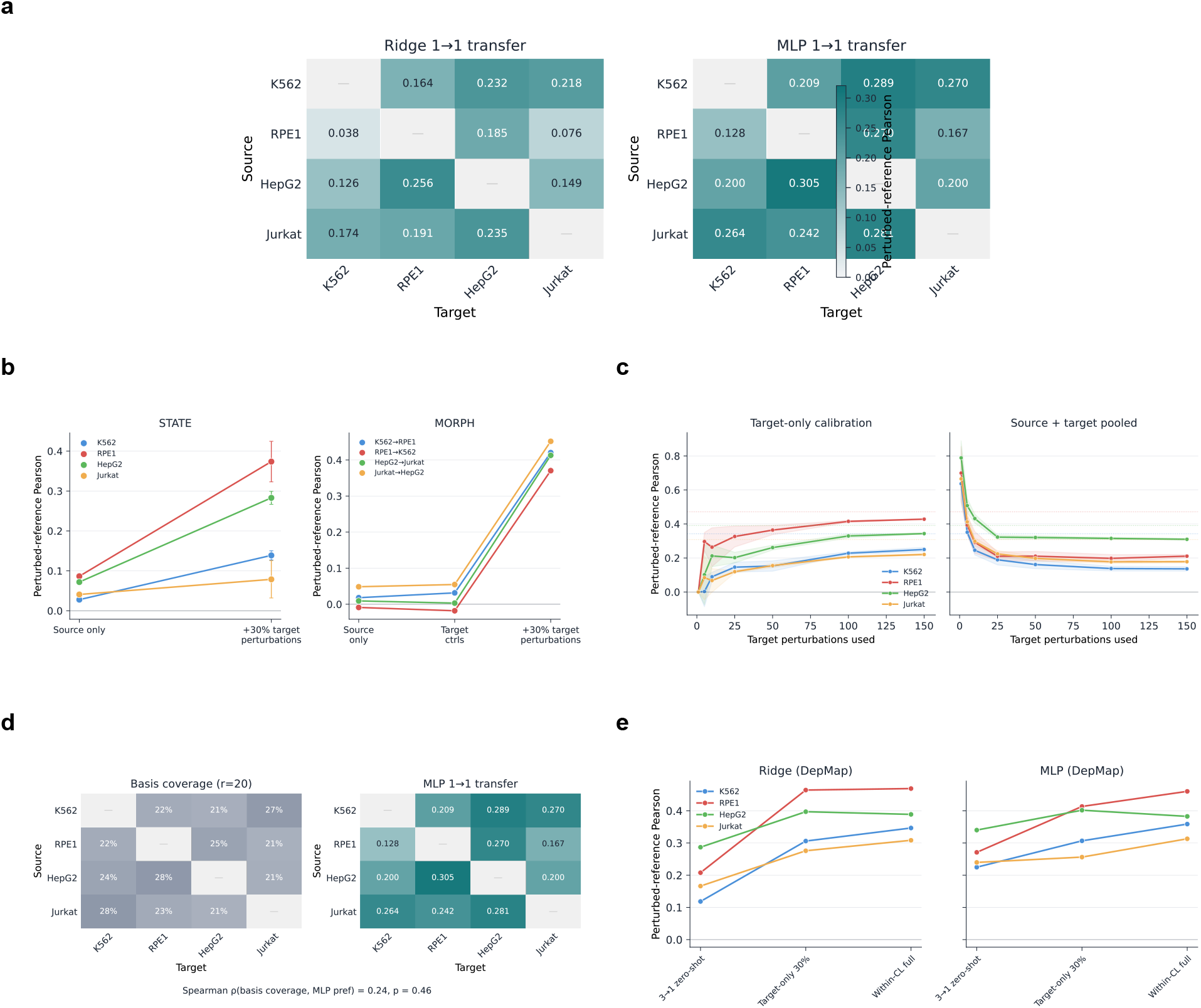
Cross cell line transfer and calibration controls. a) One source to one target transfer for DepMap based Ridge and MLP models. Rows indicate the source cell line and columns indicate the target cell line. Values show perturbed reference Pearson correlation. MLP improved over Ridge across most directed transfer pairs but performance remained target dependent. b) Effect of target information on STATE and MORPH cross cell line transfer. STATE improved when 30% target perturbations were added to training. MORPH showed little improvement from target controls alone but improved strongly when target perturbations were included. This indicates that baseline target expression is not sufficient to recover the missing target response structure. c) Target calibration saturation curves. Target only calibration improved as more target perturbations were used. Source plus target pooled training gave stronger performance with few target perturbations but showed limited additional gains as calibration size increased. Dashed lines show within cell line reference performance. d) Source target response basis coverage compared with MLP one source to one target transfer. Basis coverage was measured as the fraction of target template removed residual variance spanned by the source response basis at rank 20. Transfer performance did not track basis coverage. This indicates that DepMap based transfer does not primarily reuse source transcriptional response geometry. e) Reference baselines for DepMap based Ridge and MLP models. Performance is compared across three source to one target zero shot prediction target only training with 30% target perturbations and full within cell line training. Target perturbation measurements improved performance relative to zero shot transfer and full within cell line training provided an upper reference for matched context prediction.

**Figure 9.**
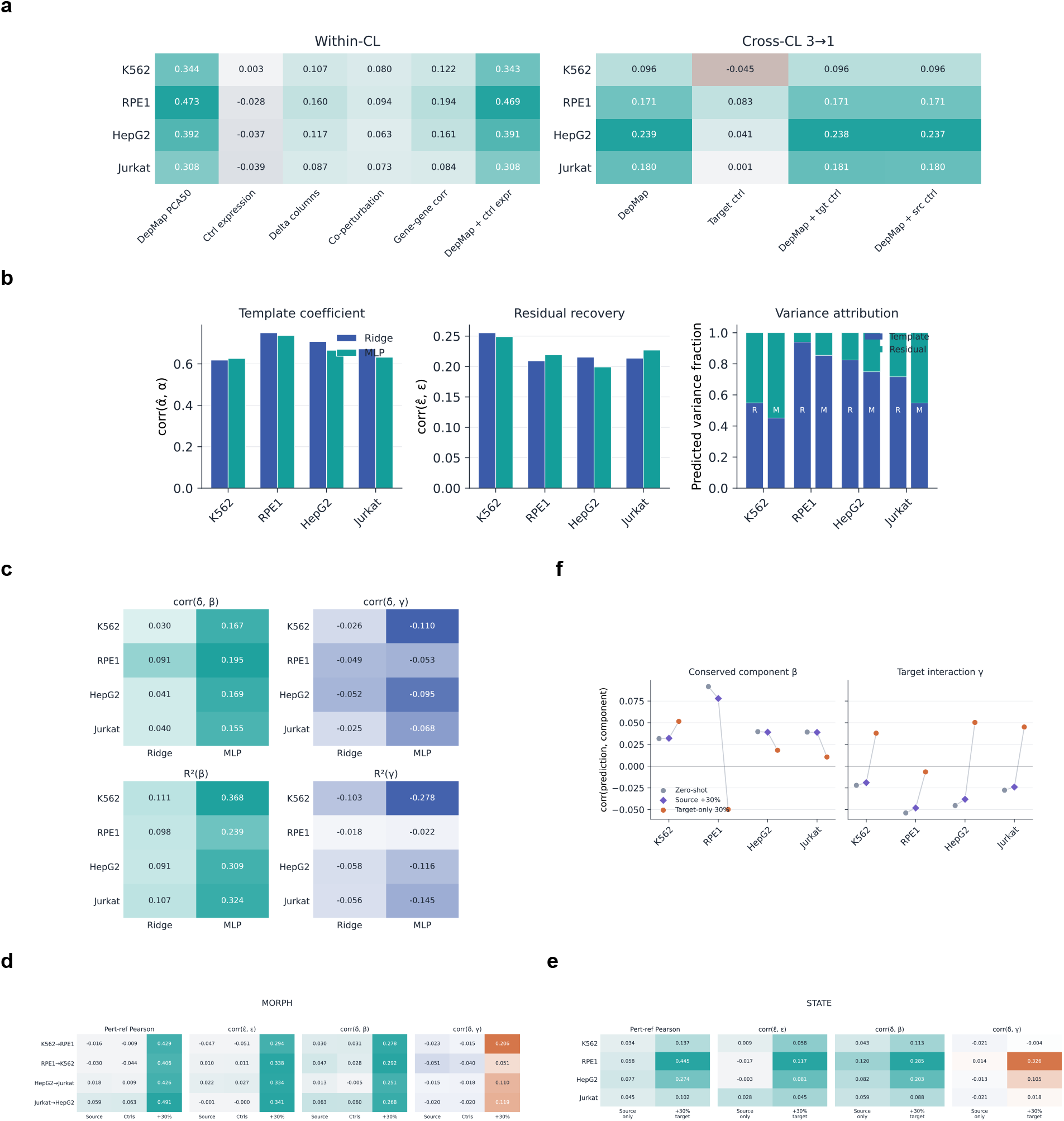
Prediction decomposition and component attribution controls. a) Descriptor and context ablation for within cell line and cross cell line prediction. Within cell line heatmaps compare DepMap PCA50 with expression derived descriptors and combined descriptors. Cross cell line heatmaps compare DepMap only target controls DepMap plus target controls and DepMap plus source controls. DepMap carried most of the predictive signal. Adding control expression descriptors did not consistently improve perturbed reference performance. b) Within cell line prediction decomposition for DepMap based Ridge and MLP models. Predicted responses were decomposed into template coefficient recovery residual recovery and predicted variance attribution. Both models recovered template coefficient variation more strongly than residual variation. Predicted variance remained substantially aligned with the shared template. c) Cross cell line zero shot attribution for DepMap based Ridge and MLP models. Predictions were compared with the conserved perturbation component *β* and the target interaction component *γ*. Models showed positive alignment with *β* but weak or negative alignment with *γ*. MLP recovered the conserved component more strongly than Ridge. d) MORPH cross cell line component attribution under source only transfer target control fine tuning and fine tuning with 30% target perturbations. Target controls alone did not recover residual or interaction signal. Adding target perturbations improved perturbed reference performance residual recovery conserved component recovery and target interaction alignment. e) STATE cross cell line component attribution under source only transfer and training with 30% target perturbations. Adding target perturbations improved perturbed reference performance and increased recovery of both conserved perturbation signal and target interaction signal. f)Target calibration attribution for DepMap based models. Source only zero shot prediction, target calibrated pooled training and target only training were compared by alignment with *β* and *γ*. Target perturbation measurements primarily increased alignment with the target interaction component *γ*, while recovery of the conserved component *β* changed less.

For generalization to unseen perturbations, DepMap-based models recovered the cell-line template more strongly than the perturbation-specific residual (Fig. 4a; Supplementary Fig. 5b). Predicted and observed template components were strongly correlated (r=0.62-0.75), whereas recovery of template-removed component was weaker but consistently positive (r=0.20–0.26). Predicted variance showed the same imbalance, with Ridge assigning 55% of its variance to the template direction in K562, 94% in RPE1, 82% in HepG2 and 72% in Jurkat. Thus, even models with positive perturbation-specific performance remained dominated by the more accessible cell-line specific component conserved across perturbations.

For generalization to unseen cell lines, attribution exposed a sharper boundary between generalizability of conserved and context-specific components. MLP predictions aligned more strongly with the perturbation-specific component conserved across cell lines than Ridge predictions (r=0.25–0.39 versus 0.10–0.13), explaining their advantage in held-out-cell-line generalization (Fig. 4b; Supplementary Fig. 5c). However, neither model recovered the cell-line-by-perturbation interaction. Whereas the state-of-the-art models, such as STATE and MORPH, generalize poorly in terms of both the perturbation-specific component and the interaction component (Fig. 4c; Supplementary Fig. 5d,e). Adding source- or target-cell-line control expression, including fine-tuning MORPH on target controls, did not materially alter this attribution (Supplementary Fig. 5a; Supplementary Note 16). The control cell state therefore is not predictive of the different perturbation response components in the held-out cellular context. Given that all benchmarked methods were unable to predict the interaction component in held-out context, we asked if this interaction component is learnable from the current perturbation data. For each target cell line, instead of a zero-shot prediction, we trained the models on 30% of all perturbations in the target cell line, with the generalization performance measured in the remaining 70% perturbations not seen in the target cell line. This improved the performance of STATE and MORPH to align positively with *γ*(*c, p, g*) (Fig. 4c; Supplementary Fig. 5d,e, Supplementary Note 15). For MORPH, this increased overall performance and the prediction of the interaction components across target cell lines, but not the perturbation-specific component conserved across cell lines (Fig. 4c-d). Importantly, comparable performance for the interaction component can be achieved when disjoint sets of perturbations in the source and target cell lines are used in training, indicating that there are learnable and generalizable patterns of cell line-by-perturbation interactions (Fig. 4d; Supplementary Fig. 5f). Thus, decomposed attribution shows that simple response-aligned models predict the conserved perturbation-specific component in unseen cell lines, and that the cell-line-by-perturbation interaction is learnable from currently available data.

These analyses reveal that models preferentially recover response components predictable from their inputs, and that this structure is invisible to aggregate scores. Response components conserved across cell lines or perturbations can be learned from baseline states, biological priors and source-cell-line responses, and simple response-aligned maps recover them as well as or better than state-of-the-art architectures. In contrast, cell-line-by-perturbation interaction is hard to predict in unseen cell lines but is learnable given the current data. This boundary persisted across linear, nonlinear and conditional generative models, indicating that architectural complexity alone cannot overcome missing target-specific information.

## 3 Discussion

Our results frame perturbation prediction as a problem of predicting distinct response components under constraints imposed by the information available to a model. By separating transcriptional responses into global, perturbation-specific, cell-line-specific, and perturbation-by-cell-line interaction components, our decomposition framework distinguishes variation only observable in the target perturbation context from variation predictable from control expression, biological priors or measured responses in training contexts. These components differ substantially in dimensionality, conservation across contexts, and predictability, indicating that they should not be treated as equivalent prediction targets. Instead, our framework identifies a hierarchy of predictability among the response components determined by the correspondence between model inputs and response structure. This component-level view provided by our framework enables identification of response components and geometry to model inputs and predictions.

The decomposition explains why aggregate metrics provide an incomplete view of perturbation-specific performance. The generic global response component is low dimensional, recurrent across perturbations, associated with proliferation arrest and cellular stress, and largely represented within control expression. Essential-gene perturbations loaded more strongly onto this component than non-essential perturbations while retaining similar underlying biological programs, suggesting that its prominence reflects the amplitude and consistency of a shared response rather than a distinct class of essential-gene biology. By contrast, the context-dependent components are higher dimensional and contain both perturbation-associated effects shared across cell lines and interactions specific to cell-line-perturbation combinations. Full-response metrics can therefore assign substantial attribution to recovery of shared structure even when perturbation-specific and context-specific effects remain less accurately predicted [12, 13]. This limits the conclusions that can be drawn from standard aggregate metrics about the prediction of perturbation identity and context dependence.

This distinction is particularly apparent in cross-cell-line prediction. Responses measured in a source cell line, together with external perturbation priors, supported prediction of part of the perturbation-specific component shared across contexts, but did not recover the perturbation-by-cell-line interaction component describing the response in the held-out cell line. Consistent with the limited correspondence between control expression and this interaction component, target control expression did not materially change this result. Whereas in our few-shot learning setup, introducing measured perturbation responses from the target cell line substantially increased interaction recovery by providing information about the response geometry of the target cell line. DepMap coessentiality profiles [14] contained transferable information about perturbation function, but their native geometry did not correspond directly to template-removed transcriptional response similarity. Predictive signal emerged after supervised alignment to response space, and relatively simple linear and non-linear mappings matched or outperformed more complex conditional models using the same prior [9–11]. At the same time, nonlinear mappings improved cross-cell-line and pathway-level recovery in several settings, indicating that additional expressivity better decodes information already present in the input. Performance therefore depends jointly on whether a prior contains information relevant to the target component and whether the decoding model can align that information with response space.

The information required to predict a perturbation component also differ across perturbation modalities. For genetic perturbations, coessentiality provides an external functional prior that supports generalization across held-out genes, cellular contexts, and previously unobserved gene combinations. For chemical perturbations, the available structure- and bioactivity-based priors did not provide an analogue with comparable predictive value. In Tahoe-100M [17], responses measured in other cell lines were substantially more predictive than Morgan fingerprints [18] or Chemical Checker signatures [19], and this transfer remained strong after removal of the generic chemical-response template. Thus, transferable perturbation representations need not take the same form across modalities. These findings have direct implications for model evaluation. Performance before and after removal of shared response structure should be considered separately, and recovery of perturbation-specific and interaction components should not be inferred from full-response metrics alone. Cross-context studies should also distinguish zero-shot transfer from settings that use measured responses in the target context, even when only a limited calibration set is available. Comparisons between architectures are most informative when models are given the same perturbation priors and target-context information, together with simple supervised mappings that establish how much of the available signal can be recovered without additional architectural complexity.

Several limitations define the scope of these conclusions. Our primary analyses use pseudobulk transcriptional responses from four CRISPRi screens enriched for essential-gene perturbations, with combinatorial and chemical perturbations serving as complementary settings. The additive decomposition is intended as a diagnostic description of response variation rather than a complete mechanistic model, and the estimated components need not correspond to uniquely separable biological processes. Gene-level pseudobulk responses also remove cell-to-cell heterogeneity that may contain additional information about perturbation state and context. Measurements of regulatory state, lineage, perturbation efficiency, temporal dynamics or richer single-cell distributions may contain context information not represented in baseline expression.

Together, these results suggest that progress in perturbation prediction should be assessed not only by how accurately a model reconstructs the complete response, but by which response components it predicts and what information makes their recovery possible. Shared response structure, perturbation-specific effects and cell-line-dependent interactions differ in their geometry, transferability and information requirements, and improvement in one component need not imply progress in the others. More broadly, treating perturbation prediction as response component estimation under predictability constraints offers a framework for designing models, datasets and benchmarks around the biological variation that a given prediction setting can support.

## 4 Methods

### 4.1 Data resources and preprocessing

#### 4.1.1 Genetic Perturb-seq screens

We analyzed four CRISPRi Perturb-seq datasets in K562, RPE1, HepG2 and Jurkat cells. K562 and RPE1 single-cell expression matrices were obtained from the original Replogle et al. data distribution [4]. HepG2 and Jurkat were obtained as processed single-cell expression matrices from Nadig et al. [5]. The four datasets were used for response decomposition, template analysis, within-cell-line prediction and cross-cell-line transfer.

Each cell contained a single guide assignment. Cells targeted by different guides against the same gene were pooled, and perturbations were represented at the target-gene level. Multi-guide cells were not present in the processed datasets. Non-targeting guides were retained as controls and were not included in the reported perturbation counts.

Perturbations represented by fewer than 30 cells were excluded uniformly across the four datasets during data construction. The reported dataset dimensions therefore correspond to the final post-filtering data. The K562 dataset contained 188,590 cells, including 10,691 non-targeting control cells, across 1,383 perturbation genes. The RPE1 dataset contained 173,737 cells, including 11,485 non-targeting control cells, across 1,499 perturbation genes. The HepG2 dataset contained 96,616 cells, including 4,976 non-targeting control cells, across 1,340 perturbation genes. The Jurkat dataset contained 184,470 cells, including 12,013 non-targeting control cells, across 1,537 perturbation genes.

#### 4.1.2 Genome-wide K562 CRISPRi screen

For the comparison of essential and non-essential perturbations, we additionally used the genome-wide K562 CRISPRi response matrix from Replogle et al. [4], distributed as `K562_gwps_normalized_bulk_01.h5ad`. This resource contains gemgroup-normalized pseudobulk perturbation profiles rather than single-cell expression values or simple control-subtracted responses.

Each row represents a gene-by-promoter perturbation condition, and each column represents a measured expression gene. Perturbation profiles were normalized by the original authors relative to non-targeting control cells from the same gemgroup. The resulting values are Z-normalized responses that account for differences in the matched control distribution across experimental batches.

The original matrix contained 11,258 gene-by-promoter conditions across 8,248 measured genes. Gene names and promoter identities were parsed from the condition index. Conditions targeting different promoters of the same gene were retained as separate perturbation profiles and therefore contributed independently to template and dimensionality analyses. Rows containing one or more non-finite values were excluded. This removed 2,341 conditions and left 8,917 finite perturbation profiles representing 7,827 unique target genes.

Essential genes were defined by membership in the focused Replogle K562 essential-gene CRISPRi library rather than by an external essentiality annotation. The finite genome-wide matrix contained 1,046 unique essential-gene targets represented by 1,056 promoter-specific perturbation conditions. Essential perturbations were not required to pass the energy-test significance threshold for the initial library-based comparison. For significance-matched analyses, both essential and non-essential perturbations were restricted to conditions with energy-test *p <* 0.001, yielding 894 essential and 1,574 non-essential perturbation conditions.

#### 4.1.3 Combinatorial CRISPRa benchmark

For combinatorial perturbation prediction, we used the Norman CRISPRa Perturb-seq benchmark [15]. The dataset contains K562 cells profiled under single-gene perturbations, double-gene perturbations and control conditions. Expression was evaluated in a 2,000-gene space, and pseudobulk responses and perturbed-reference Pearson correlations were calculated as described below.

The benchmark split contained 137 training conditions, comprising 70 single-gene perturbations, 36 double-gene perturbations and 31 control-paired single-gene conditions. The test set contained 116 conditions, comprising 37 single-gene and 79 double-gene perturbations. Unseen-gene stratification, additive baselines and DepMap pair-feature models are described in the combinatorial perturbation prediction section.

#### 4.1.4 Chemical perturbation benchmark

For chemical perturbation transfer, we used the Tahoe-100M dataset [17]. The analysis included C32, HOP62, HepG2/C3A, Hs 766T and PANC-1 cells profiled across 268 compounds at doses of 0.05, 0.5 and 5.0 *µ*M. DMSO-treated cells were used as controls.

Pseudobulk drug responses were calculated as the difference between the mean expression of compound-treated cells and the corresponding DMSO control expression. Unless otherwise stated, analyses used 2,000 highly variable genes and required at least 20 cells per cell-line-by-compound-by-dose condition.

Chemical perturbation analyses compared static compound descriptors with source-response representations derived from measured responses to the same compound in other cell lines. Source-response transfer, held-out-compound evaluation, leave-one-mechanism-of-action-out evaluation, template-centering controls and compound-label permutation controls are described in the chemical perturbation transfer section.

#### 4.1.5 Gene essentiality descriptors

DepMap dependency profiles [14] were used as biological perturbation descriptors for Ridge and MLP genetic perturbation models. We used the `CRISPRGeneEffect` matrix, containing dependency scores for 18,531 genes across 1,208 DepMap cell lines. For each perturbation target, the corresponding gene-dependency vector was used when available. Perturbation genes without a matching DepMap profile were excluded from DepMap-based analyses.

For supervised prediction, principal component analysis was fitted using training-set DepMap profiles and subsequently applied to both training and held-out perturbations. Unless otherwise stated, the first 50 principal components were used as the perturbation descriptor. To determine how predictive information was distributed across the DepMap eigenspectrum, we additionally evaluated ranks of 1, 5, 10, 20, 50, 100 and 200, together with the complete unreduced dependency profile.

#### 4.1.6 Chemical descriptors

For chemical perturbation analyses, compounds were matched to InChIKeys using an existing drug-name mapping, with salt, hydrate and solvate suffixes removed when required for name matching. Canonical SMILES were retrieved from PubChem using the matched InChIKey or, when unavailable, the cleaned compound name. SMILES were parsed using RDKit, and compounds that could be successfully parsed were represented by 2,048-bit radius-2 Morgan fingerprints [18].

Chemical Checker representations were generated from the canonical SMILES using the `signaturizer` package [19]. We included the non-transcriptomic Chemical Checker spaces A1–A5, B1–B5, C1–C5 and E1–E5, corresponding to chemical structure, target and bioactivity information, biological networks and processes, and cell-based phenotypes, respectively. Transcriptomic D spaces were excluded to prevent leakage from transcriptional response measurements. The 128-dimensional signatures produced for each available space were concatenated to obtain a single Chemical Checker representation for each compound; missing or invalid signatures were omitted from the concatenation.

#### 4.1.7 Template and pathway-level analyses

Hallmark gene sets were obtained from the 2024.1 human release of MSigDB [20]. Gene symbols were matched to the highly variable genes used in each analysis, with Ensembl identifiers mapped to gene symbols where necessary. Pathways represented by fewer than five genes in the corresponding expression space were excluded.

For pathway-level prediction analyses, each retained gene set was represented as an equal-weight binary vector over the expression feature space and normalized to unit Euclidean norm. For each perturbation, pathway activity was calculated by projecting the template-removed transcriptional response onto this vector. Prediction performance for each pathway was then quantified as the Pearson correlation between the predicted and observed pathway activities across held-out perturbations. For models evaluated by five-fold cross-validation, pathway activities were pooled across folds before computing the correlation. For models evaluated across multiple random seeds, predictions were aggregated across the available seeds before pathway-level performance was calculated.

#### 4.1.8 Expression preprocessing and gene selection

Gene symbols were used as expression-feature identifiers throughout the four-cell-line genetic perturbation analysis. The gene spaces of K562, RPE1, HepG2 and Jurkat were intersected, yielding 6,642 genes shared across all four datasets. No missing or duplicated gene symbols were present after intersection. Mitochondrial genes were retained.

Within each cell line, expression values were normalized to a total count of 10,000 per cell using `scanpy.pp.normalize_total` and transformed using log(1 + *x*). The resulting expression matrices were concatenated across cell lines without balancing or downsampling the number of cells contributed by each cell line.

Two thousand highly variable genes were selected once from the concatenated expression matrix using `scanpy.pp.highly_variable-genes` with `flavor=“seurat”` and without supplying a batch or cell-line key. This produced a single highly variable gene set based on expression variation across the pooled cells. The same 2,000 genes, retained in the same order, were used for all four cell lines and for analyses requiring direct comparison across cell lines.

No gene-wise scaling was applied after normalization and log transformation. Pseudobulk expression profiles were therefore calculated directly from log-normalized, unscaled expression. No additional cell-level quality control or batch correction was applied beyond the filtering provided in the source datasets.

### 4.2 Response construction and evaluation

#### 4.2.1 Pseudobulk perturbation responses

For each perturbation *p* in cell line *c*, cells were grouped using the gene-level perturbation annotation. Cells carrying different guides targeting the same gene were therefore pooled before pseudobulk construction. Pseudobulk perturbation responses were computed directly from log-normalized, unscaled expression by subtracting the mean expression of non-targeting control cells from the mean expression of cells assigned to each perturbation:

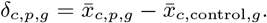

Here, *g* denotes the measured output gene. Perturbations represented by fewer than 30 cells were excluded uniformly across K562, RPE1, HepG2 and Jurkat before pseudobulk construction. Analyses involving multiple cell lines were performed on the shared perturbation and gene sets defined for the corresponding analysis.

The same pseudobulk response definition was used for within-cell-line prediction, cross-cell-line transfer, combinatorial perturbation prediction and chemical perturbation transfer, with dataset-specific controls and filtering described in the corresponding sections.

#### 4.2.2 Perturbation-level splits

For within-cell-line prediction analyses, we used perturbation-level train and test splits. All cells assigned to the same gene-level perturbation were kept in the same split. Unless otherwise stated, 85% of perturbations were used for training and 15% for testing, with random seed 42. For matched within-cell-line comparisons of Ridge, MLP and MORPH, we used five-fold perturbation-level cross-validation restricted to perturbations with available DepMap embeddings. All cells assigned to the same gene-level perturbation were kept in the same fold, and the same held-out perturbations were used for each model within a fold.

For target-calibrated cross-cell-line analyses, target perturbations were split into calibration and test sets. Unless otherwise stated, 30% of target perturbations were used for calibration and the remaining 70% were held out for evaluation. The same target calibration and test split was used across models.

For zero-shot cross-cell-line analyses, no perturbation responses from the target cell line were used during training. For double-unseen analyses, both the target cell line and a subset of perturbation genes were held out during training. Task-specific details for one-source, three-source, target-calibrated and double-unseen transfer are described in the cross-cell-line prediction section.

#### 4.2.3 Perturbed-reference Pearson correlation

For each cell line *c*, the training perturbed centroid was defined as

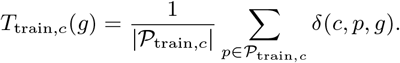

Standard Pearson correlation was computed between the predicted and true pseudobulk responses:

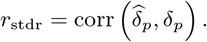

Perturbed-reference Pearson correlation was computed after subtracting the training perturbed centroid from both the prediction and the ground truth [13]:

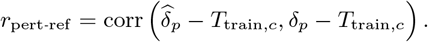

This metric measures prediction of perturbation-specific response variation relative to the training perturbation centroid. Unless otherwise stated, model performance was summarized as the mean perturbed-reference Pearson correlation across held-out perturbations.

### 4.3 DepMap response mapping and within-cell-line prediction

#### 4.3.1 DepMap perturbation descriptors

For each perturbation target gene, we used its DepMap dependency profile [14] as an external perturbation descriptor. Each profile was represented as a vector of gene-effect scores across 1,208 DepMap cell lines. DepMap-based analyses were restricted to perturbation genes with a matching dependency profile.

For supervised prediction, perturbation genes were treated as observations and DepMap cell lines as descriptor features. Principal component analysis was fitted using the training perturbations within each fold and subsequently applied to both training and held-out perturbations. Unless otherwise stated, the first 50 principal components were used as the perturbation descriptor.

#### 4.3.2 DepMap similarity and perturbation-response coherence

To test whether proximity in DepMap space corresponded to similarity in transcriptional response space, we compared pairwise similarities among perturbation genes. Within each cell line, cosine similarity was calculated between the raw, unreduced DepMap dependency profiles of all eligible perturbation-gene pairs. For the same pairs, cosine similarity was calculated between their full pseudobulk responses and between their template-orthogonal residual responses.

For perturbation *p*, the template-orthogonal residual was defined as

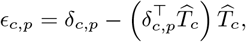

where 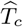 denotes the unit-normalized template direction for cell line *c*. The association between DepMap similarity and response similarity was quantified using the Spearman correlation between the upper-triangular entries of the corresponding pairwise cosine-similarity matrices. Correlations were calculated separately for full responses and template-orthogonal residuals, allowing us to assess whether DepMap similarity captured perturbation-specific response structure beyond the shared template.

We performed two complementary analyses of local response coherence in DepMap space. First, gene pairs were divided into deciles according to their DepMap cosine similarity, and the mean full-response and template-orthogonal residual similarity was calculated within each decile.

Second, perturbation genes were clustered from their raw 1,208-dimensional DepMap dependency profiles using *k*-means, without PCA reduction or feature standardization. We evaluated

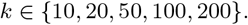

Clustering was performed using `scikit-learn KMeans` with `random_state=0` and `n_init=5`. For each value of *k*, the mean template-orthogonal residual similarity was calculated separately for gene pairs within the same cluster and for pairs assigned to different clusters. Between-cluster pairs were subsampled for computational efficiency. All analyses were performed independently within each cell line.

#### 4.3.3 Within-cell-line response prediction

Ridge regression, multilayer perceptron and MORPH models were trained to predict the full pseudobulk response of a perturbation from its DepMap descriptor. A separate model was fitted for each cell line:

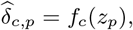

where *z*_*p*_ denotes the PCA-reduced DepMap profile of perturbation gene *p*, and 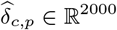 is its predicted transcriptional response in cell line *c*.

Models were compared using five-fold perturbation-level cross-validation restricted to perturbation genes with available DepMap profiles. Principal component analysis was refitted within each training fold. Ridge, MLP and MORPH were trained and evaluated using the same held-out perturbations in each fold. Performance was measured using perturbed-reference Pearson correlation, with the reference centroid calculated from the training perturbations of the corresponding fold.

Ridge regression was fitted as a multi-output regression using `scikit-learn`, with regularization parameter *α* = 10. The same regularization setting was used across cell lines and cross-validation folds.

The MLP contained two hidden layers of 256 units with ReLU activations [21] and mapped the DepMap PCA representation to the 2,000-gene response vector. The model was optimized using mean-squared-error loss and Adam [22], with an initial learning rate of 5 *×* 10^−5^, weight decay of 10^−5^, mini-batches of 64 and gradient clipping at a norm of 1.0. Models were trained for a fixed 400 epochs using a cosine-annealing learning-rate schedule, without early stopping or a separate validation set. A single random seed, equal to the cross-validation fold index, was used in each fold. The same architecture and training configuration were used for every cell line, and no hyperparameters were tuned separately by fold or cell line.

MORPH was trained and evaluated using the same perturbation folds and DepMap descriptors, as described in the model-specific section below.

#### 4.3.4 DepMap PCA rank ablation

To determine whether predictive information was concentrated in the leading DepMap axes or distributed across lower-variance dimensions, we repeated the Ridge and MLP analyses using

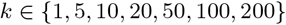

principal components. We additionally evaluated the complete unreduced DepMap dependency profile. For each representation rank, PCA was fitted using the training perturbations within each fold, while the perturbation splits, model configurations and evaluation metric were held fixed.

### 4.4 Model-specific perturbation predictors

#### 4.4.1 scGPT fine-tuning and inference

We evaluated scGPT [9] as a pretrained foundation-model baseline for within-cell-line perturbation prediction. scGPT was initialized from the `scGPT_human` whole-human pretrained checkpoint distributed by the model authors and fine-tuned separately for each cell line. Architecture hyperparameters were inherited from the checkpoint configuration, and data were loaded using the GEARS `PertData` pipeline [6].

scGPT was evaluated using a single perturbation-level split containing approximately 85% training and 15% test perturbations. This split was replicated across five random seeds and differed from the five-fold cross-validation used for Ridge, MLP and MORPH. Training perturbations were further divided at the perturbation level into fitting and validation subsets, ensuring that cells assigned to the same perturbation remained in the same subset.

The pretrained gene encoder, value encoder and transformer encoder weights were loaded and updated during full fine-tuning. No pretrained layers were frozen. The expression decoder was initialized from scratch and implemented as an MLP rather than the default affine decoder, thereby avoiding an explicit skip connection from control expression to predicted expression.

Fine-tuning minimized mean-squared error between predicted and observed post-perturbation expression across gene positions. Adam [22] was used with a learning rate of (1 *×* 10^−4^), a `StepLR` schedule with a multiplicative decay of 0.9 per epoch, gradient clipping at a norm of (1.0), a batch size of 64 and mixed-precision training. No auxiliary scGPT objectives were used. Training was performed for up to 15 epochs, with early stopping based on validation mean-squared error and a patience of five epochs. The checkpoint with the lowest validation loss was used for inference.

During training, gene positions were randomly subsampled to a maximum sequence length of 1,536 per batch to satisfy memory constraints. At inference, the complete shared gene set was processed without subsampling, and predictions were extracted for the 2,000 highly variable evaluation genes.

For each held-out perturbation, all available non-targeting control cells from the corresponding cell line were paired with the perturbation identity and processed in batches of 64. Predicted post-perturbation expression profiles were averaged across control cells to obtain a pseudobulk prediction. The same control-cell pool was used for all perturbations and seeds. Predicted perturbation responses were calculated by subtracting the non-targeting control centroid and evaluated using perturbed-reference Pearson correlation.

#### 4.4.2 MORPH training and inference

MORPH [11] was trained using the original implementation, with DepMap dependency profiles supplied as perturbation descriptors. For within-cell-line prediction, MORPH was trained separately in each cell line for up to 100 epochs using Adam [22] with a learning rate of (1 *×* 10^−3^), a batch size of 32, an MMD kernel bandwidth of (*σ*=1500), (*γ*_1_=1.0) and mean-squared-error reconstruction loss. Early stopping with a patience of 20 epochs monitored the total validation loss on perturbation-level held-out validation data, and the checkpoint with the lowest validation loss was restored for inference. Five random seeds were used for each cell line.

For source-only cross-cell-line transfer, MORPH was trained in the source cell line and applied directly to target-cell-line control cells. For each test perturbation, 100 non-targeting target cells were sampled with replacement, paired with the corresponding DepMap descriptor and passed through the model. Predicted single-cell expression profiles were averaged, and the predicted response was calculated by subtracting the target control centroid. Five random seeds were used for each directed source-to-target pair.

For target-control fine-tuning, all MORPH parameters were updated for nine epochs using target non-targeting cells. Individual control cells were passed through the encoder-decoder with the null perturbation embedding. The MMD term was retained, and reconstruction loss was combined with a linearly annealed KL-divergence term whose weight increased from 0 to 1 over the nine epochs. Fine-tuning used Adam at (1 *×* 10^−3^) and gradient-value clipping at 0.5. All target control cells were used, and no validation split or early stopping was applied during this fixed fine-tuning schedule.

For target-perturbation calibration, fine-tuning additionally included supervised batches from the target calibration perturbations. Each epoch alternated between target-control reconstruction batches and perturbation-prediction batches. Prediction batches paired target control cells with the corresponding DepMap descriptor and were supervised against the observed target pseudobulk expression profile.

For combinatorial perturbation prediction, MORPH was trained using the original implementation and data pipeline with the common Norman benchmark split. Single perturbations were encoded through the perturbation encoder. For double perturbations, the two DepMap profiles were encoded independently and their latent representations were summed after the perturbation encoder and before the cross-attention mechanism. Training used the same optimizer and loss settings, with up to 100 epochs, batch size 32, early stopping with patience 20 and three random seeds. At inference, 100 control cells were sampled for each condition, predicted expression profiles were averaged, and the control centroid was subtracted to obtain the predicted response.

#### 4.4.3 Linear probing of MORPH representations

To quantify how response information changed along the MORPH perturbation pathway, we extracted representations at five stages: the raw DepMap profile, the output of the perturbation encoder, the representation obtained after combining perturbation and cell-context inputs but before cross-attention, and the outputs of the first and second post-attention layers.

MORPH parameters were frozen during representation extraction. For the cell-context-dependent stages, representations were averaged across 100 randomly sampled control cells to obtain one feature vector per perturbation. The same five-fold perturbation splits used for the MORPH prediction analysis were used for linear probing.

Within each fold, intermediate representations were reduced using PCA fitted on the training perturbations. We retained at most 50 components, or (*P*_train_ − 1) components when fewer training perturbations were available. A Ridge regression model with (*α*=10) was then fitted from each representation to the pseudobulk perturbation response and evaluated on the corresponding held-out perturbations using perturbed-reference Pearson correlation. Every representation stage was evaluated using the same training and test perturbations.

This analysis quantified how much held-out, centroid-relative response information was linearly recoverable from each stage of the frozen MORPH perturbation pathway.

#### 4.4.4 STATE training and inference

STATE [10] was trained using the original implementation with a GPT-2 transformer backbone [23]. Training was performed for 80,000 gradient steps using a batch size of 8, a learning rate of (1 *×* 10^−3^), a hidden dimension of 128, four encoder layers, four decoder layers, eight attention heads, eight key-value heads, a cell-set length of 32 and residual prediction. The model was optimized using its distributional training objective, which compares predicted and observed populations of perturbed cells without requiring one-to-one correspondence between individual control and perturbed cells.

Perturbations were represented using one-hot encodings constructed from the perturbation labels present in the training data. For zero-shot transfer, the vocabulary therefore contained only source-cell-line perturbations. For calibration experiments, it was expanded to include the target calibration perturbations. Perturbations absent from the training vocabulary could not be encoded and were excluded from STATE double-unseen evaluation.

Checkpoints were saved every 2,000 steps and assessed using a perturbation-level validation set disjoint from both training and test perturbations. The selected validation checkpoint was used for inference. Five training seeds were evaluated for each configuration.

At inference, 32 non-targeting control cells from the target cell line were sampled and supplied as the cellular-context set together with each test perturbation identity. STATE generated a predicted post-perturbation profile for each input cell. Predictions were averaged across the cell set to obtain a pseudobulk expression profile, and the target control centroid was subtracted to obtain the predicted perturbation response. Random seeds varied model initialization, training randomness and control-cell sampling.

For target calibration, 30% of eligible target perturbations and their observed expression profiles were added to the training data. The remaining 70% were withheld for evaluation.

### 4.5 Cross-cell-line prediction tasks

#### 4.5.1 One-source and three-source cell line tasks

We evaluated cross-cell-line prediction using either one or three source cell lines. In both settings, no target-cell-line perturbation responses were available during training. Source training included all perturbations in the processed datasets with an available DepMap profile; all retained perturbations were represented by at least 30 cells.

In the one-source setting, a model was trained using perturbations from one source cell line and evaluated in one paired target cell line. We evaluated four directed transfers: K562 to RPE1, RPE1 to K562, HepG2 to Jurkat and Jurkat to HepG2.

In the three-source setting, perturbation data from the three non-target cell lines were pooled and the fourth cell line was held out entirely. Each source-cell-line-by-perturbation pair was treated as a separate training observation. A gene perturbed in multiple source cell lines therefore contributed repeated DepMap descriptors paired with different cell-line-specific responses. Ridge and MLP received no explicit cell-line descriptor and consequently estimated a source-averaged response mapping for perturbations represented in multiple source contexts.

For models using continuous DepMap descriptors, target evaluation included all target perturbations with available DepMap profiles, regardless of whether the corresponding gene was perturbed in a source cell line. The evaluation set therefore contained both context-unseen perturbations observed in at least one source cell line and perturbations absent from all source response data. STATE evaluation was restricted to target perturbations represented in its source-derived one-hot vocabulary.

For Ridge and MLP, PCA with 50 components was fitted using source-training DepMap profiles and applied to target perturbations. Zero-shot perturbed-reference evaluation used the observed target perturbation centroid as an evaluation-only reference. This centroid was used only after inference and was not available during model fitting or checkpoint selection.

#### 4.5.2 Perturbation and cell line held-out prediction

In the double-unseen setting, both the target cell line and a subset of perturbation genes were held out during training. We first identified perturbations shared between the source cell lines and the target cell line. From these shared perturbations, 30% were selected by a seeded random permutation and defined as double-unseen perturbations. These perturbations were removed from the source training data in all source cell lines, and models were retrained on the remaining source data.

Predictions were evaluated on the double-unseen perturbations in the target cell line. In this setting, the model had never observed perturbation responses for the held-out genes in any source cell line and had never observed perturbation responses in the target cell line. For comparison, we also evaluated target perturbations that remained present in the source training data. PCA was refitted on the pruned source data for each split. This setting separated the difficulty of predicting in a new cellular context from the additional difficulty of predicting perturbation genes never observed in any cell line.

#### 4.5.3 Target cell line calibration and target cell line-only reference

In the three-source-to-one-target calibration setting, 30% of target-cell-line perturbations were randomly selected as a calibration set and added to the source training data. Models were evaluated on the remaining 70% of target perturbations. The same target calibration and test split was used for all models in this setting.

For models using PCA, the projection was fitted on the combined source and target-calibration DepMap embeddings. Perturbed-reference evaluation used the target-calibration centroid as the reference vector. For STATE, the calibration perturbations were included in the one-hot perturbation vocabulary and the model was trained on data from all four cell lines, with the remaining 70% of target perturbations withheld for evaluation. For MORPH, calibration perturbations were included during fine-tuning together with target controls, using reconstruction loss on control cells and supervised prediction loss on calibration perturbations.

In the target-only calibration reference setting, models were trained only on the 30% target calibration set, without source-cell-line data. The same remaining 70% of target perturbations were used for evaluation. This setting provided a reference for prediction when target perturbation measurements were available but no source-cell-line responses were used.

#### 4.5.4 Source-target cell line response basis coverage

To test whether cross-cell-line transfer performance could be explained by overlap between source and target response subspaces, we measured the fraction of target residual-response variance captured by the source response basis. This analysis was used as a diagnostic of source-to-target response geometry and was not used during model training.

Within each cell line, the perturbation template *T* was computed as the mean response across all perturbations. The unit-normalized template direction was defined as

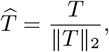

and the template-removed residual for perturbation *p* was computed as

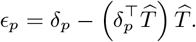

For each directed source-to-target pair, singular value decomposition was applied to the source residual-response matrix *E*_source_, with perturbations as rows and genes as columns:

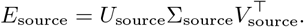

The right singular vectors in *V*_source_ defined the source response basis. Target residual responses were projected onto the first *k* source basis vectors, with

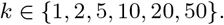

Let *V*_source,*k*_ denote the matrix containing the first *k* right singular vectors. The corresponding orthogonal projection matrix was

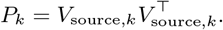

Coverage was computed as the fraction of total target residual-response variance retained after projection onto the source basis:

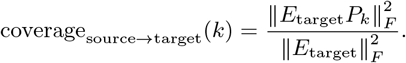

The response-basis gap was defined as

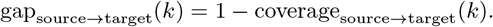

This quantity measures the fraction of target residual-response variance that is not spanned by the source response basis at rank *k*. Template removal was performed before computing basis coverage so that the analysis measured overlap in perturbation-specific variation rather than shared template structure.

### 4.6 Combinatorial perturbation prediction benchmark

#### 4.6.1 Benchmark split and unseen-gene strata

We evaluated combinatorial perturbation prediction using the Norman CRISPRa Perturb-seq benchmark [15]. Expression was represented in a Norman-specific 2,000-gene space selected by the GEARS preprocessing pipeline from the Norman K562 expression matrix. This evaluation space differs from the shared 2,000-gene set used for the four-cell-line CRISPRi analyses. Pseudobulk responses and perturbed-reference Pearson correlations were calculated as described above. The perturbed-reference centroid was computed from all 137 training conditions.

The training split contained 101 control-paired single-gene conditions and 36 double-gene conditions. The single-gene conditions represented 70 unique genes: 70 appeared in the GENE + control orientation, and 31 of these genes additionally appeared in the reverse control + GENE orientation. Cells from both orientations targeting the same gene were pooled to obtain a single gene-level pseudobulk response.

The test split contained 37 single-gene and 79 double-gene conditions. The combinatorial analysis was restricted to the 79 test doubles; test single perturbations were not evaluated.

Test doubles were stratified according to the availability of their constituent single-gene responses in the training split. A gene was classified as observed only when its corresponding single-gene condition was present in training. Appearance solely as a constituent of a training double did not count as single-gene exposure. This produced three strata: 0*/*2 unseen, in which both constituent single-gene responses were observed; 1*/*2 unseen, in which one was observed; and 2*/*2 unseen, in which neither was observed. Before DepMap filtering, the three strata contained 14, 48 and 17 test doubles, respectively.

#### 4.6.2 DepMap coverage and evaluation sets

For each Norman perturbation gene, we extracted its dependency profile from the DepMap CRISPRGeneEffect matrix. Of the 105 perturbation genes in the benchmark, 99 had a matching DepMap profile. Missing entries within available dependency profiles were replaced with zero before feature construction.

Conditions involving any of the six genes absent from DepMap were excluded from DepMap-based analyses. This left 72 evaluable test doubles: 13 in the 0*/*2 unseen stratum, 44 in the 1*/*2 unseen stratum and 15 in the 2*/*2 unseen stratum. Ridge, MLP and MORPH were evaluated on this common set of 72 test doubles.

#### 4.6.3 Observed-single additive baseline

For test doubles in the 0*/*2 unseen stratum, we evaluated SALT [16], an additive baseline that predicts the double-perturbation response as the sum of the two measured single-gene responses, 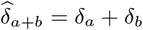, where *δ*_*a*_ and *δ*_*b*_ are the training-set pseudobulk responses for the corresponding single-gene perturbations. Because this baseline requires both constituent single-gene responses, it was evaluated only on the 13 DepMap-covered test doubles in the 0*/*2 unseen stratum.

An additive-plus-residual model was not evaluated because no training double had both constituent single-gene responses available in the training split. Consequently, the residual interaction term could not be constructed for any training example.

#### 4.6.4 DepMap pair-feature models

Ridge and MLP models using DepMap pair features did not require measured single-gene responses and were therefore evaluated across all three unseen-gene strata.

Let *v*_*a*_ and *v*_*b*_ denote the raw DepMap dependency profiles of genes *a* and *b*. For each double perturbation, we constructed the order-invariant pair representation *x*_*a*+*b*_ = *v*_*a*_ + *v*_*b*_.

Ridge and MLP readouts were trained only on the 36 training double perturbations, after excluding conditions involving genes without DepMap coverage, and mapped the summed pair descriptor to the 2,000-gene pseudobulk response. Principal component analysis was fitted using the filtered training doubles and retained at most 50 components or *n*_train_ − 1, whichever was smaller.

Ridge regularization and MLP architecture followed the settings used for the single-gene DepMap response models. The MLP contained two hidden layers of 256 units with ReLU activations and was trained for 400 epochs using cosine learning-rate annealing, weight decay of 10^−5^ and gradient clipping. Predictions were averaged across three random seeds to reduce initialization variance in the small training set.

#### 4.6.5 MORPH latent composition

MORPH was trained using the same Norman benchmark split and DepMap-covered conditions. Each constituent DepMap profile was encoded independently by the perturbation encoder. For double perturbations, the resulting latent perturbation vectors were summed after the perturbation encoder and before the cross-attention mechanism. Training and inference otherwise followed the MORPH settings described above.

#### 4.6.6 Evaluation

All combinatorial models were evaluated on the same set of test double perturbations with DepMap coverage whenever the model was defined for the corresponding stratum. Performance was summarized separately for the 0*/*2, 1*/*2, and 2*/*2 unseen-gene strata using perturbed-reference Pearson correlation, with the reference centroid computed from training perturbations only. This evaluation separates performance driven by reuse of observed single-gene responses from performance requiring generalization to double perturbations whose constituent genes were not observed as single perturbations during training.

### 4.7 Chemical perturbation transfer in Tahoe-100M

#### 4.7.1 Tahoe response construction

We evaluated chemical perturbation transfer using the Tahoe-100M dataset [17]. The analysis included C32, HOP62, HepG2/C3A, Hs 766T and PANC-1 cells profiled across 268 compounds at concentrations of 0.05, 0.5 and 5.0 *µ*M. Each compound-dose pair was treated as a distinct perturbation condition.

Two thousand highly variable genes were selected from the Tahoe expression data and used as a shared evaluation space across all five cell lines. Conditions represented by fewer than 20 cells were excluded. Of the 4,020 possible cell-line-by-compound-by-dose conditions, 3,662 passed this threshold.

For each cell line, all DMSO-treated cells were pooled to obtain a single control centroid. The response of compound-dose condition *p* in cell line *c* was defined as

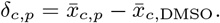

DMSO controls were not matched by plate or experimental batch. Cross-cell-line analyses used matched compound-dose conditions. When a condition was unavailable in one source cell line, its source representation was calculated from the remaining source cell lines. Fewer than 0.2% of target conditions had incomplete source coverage.

#### 4.7.2 Static compound descriptors

We evaluated whether target-cell-line responses could be predicted from static chemical identity alone. Canonical compound SMILES were converted to binary 2,048-bit radius-2 Morgan fingerprints using `RDKit` [18]. Chemical Checker signatures [19] provided an additional compound representation.

We used 20 non-transcriptomic Chemical Checker spaces, each represented by a 128-dimensional signature. These comprised spaces A1-A5, B1-B5, C1-C5 and E1-E5, which were concatenated into a 2,560-dimensional descriptor. Transcriptomic D-level spaces were excluded to prevent leakage from gene-expression measurements. Chemical Checker signatures were available for 265 of the 268 compounds.

Static descriptors were indexed by compound identity and did not include dose as an input feature. The same Morgan or Chemical Checker representation was therefore assigned to all three concentrations of each compound.

Morgan and Chemical Checker models were trained separately within each target cell line using compound-level cross-validation. Ridge and MLP models mapped the static descriptor to the target-cell-line pseudobulk response and did not use responses measured in source cell lines. These models provided a baseline for response prediction from chemical identity alone. Dimensionality reduction, when used, was fitted using training compounds only.

#### 4.7.3 Direct and low-rank source-response transfer

We next evaluated whether measured drug responses in other cell lines provided more informative perturbation representations than static chemical descriptors. For target cell line *t* and compound–dose condition *p*, the source-response vector was defined as the element-wise average response across the remaining cell lines in which that condition was available:

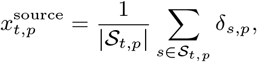

where *S*_*t,p*_ denotes the set of available source cell lines for condition *p*.

As a direct-transfer baseline, the averaged source-response vector was used as the target prediction:

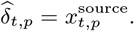

This baseline tests whether the measured transcriptional response of a compound transfers directly across cellular contexts without learning a target-specific mapping.

We additionally trained low-rank source-response Ridge models. For each target cell line, the training source-response vectors were arranged as rows of a matrix *X*_source_. Singular value decomposition was fitted using training compounds only:

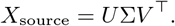

The first *k* right singular vectors defined a source-response program basis, and each compound–dose condition was represented by its coordinates in this basis. We evaluated

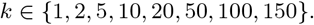

A Ridge model with *α* = 10 mapped the source-response coordinates to the target-cell-line pseudobulk response. Both the singular-vector basis and the Ridge mapping were fitted using training compounds and subsequently applied to held-out compounds.

#### 4.7.4 Held-out-compound and leave-one-MoA-out evaluation

We evaluated generalization using held-out-compound and leave-one-mechanism-of-action-out settings.

For held-out-compound evaluation, unique compounds were randomly partitioned into five folds using seed 42. All doses of a compound were assigned to the same fold to prevent dose-level leakage. The same fold assignments were used across target cell lines and models.

For leave-one-mechanism-of-action-out evaluation, all compounds assigned to one annotated mechanism were held out together. Mechanism-of-action annotations were available for 113 compounds. Compounds annotated as unclear and mechanism classes containing fewer than three compounds were excluded. For compounds with multiple annotations, the first listed mechanism was used as the primary label.

Nine mechanism classes comprising 37 compounds were evaluated: DNA synthesis or repair inhibitors, cyclooxygenase inhibitors, EGFR or ERBB inhibitors, JAK or STAT inhibitors, glucose transporter inhibitors, microtubule inhibitors, CDK inhibitors, RAF inhibitors and MTOR inhibitors.

Performance was calculated as the Pearson correlation across genes between predicted and observed pseudobulk responses for each held-out compound-dose condition and then averaged across conditions. Centroid-relative performance was calculated after subtracting the target training-response centroid from both the prediction and observation.

#### 4.7.5 Centroid-centering and permutation controls

To determine how much source-response transfer depended on a shared drug-response centroid, we repeated the low-rank analysis after centering source and target responses.

For each source cell line *s*, the training centroid was defined as

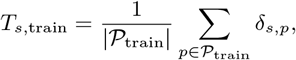

and source responses were centered as

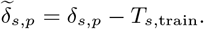

The centered source representation was obtained by averaging 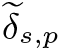 across the available source cell lines. Target responses were centered using the corresponding target training centroid, *T*_*t*,train_. The SVD basis and Ridge model were then fitted in this centroid-relative response space.

To test whether transfer depended on matching the same compound-dose condition across source and target cell lines, we performed a label-permutation control. Source-response vectors were randomly shuffled across compound-dose labels while target responses and train-test partitions were left unchanged. We performed 100 permutations using sequential random seeds from 0 to 99. The resulting null distribution was compared with performance obtained using the correctly matched source-response labels.

#### 4.7.6 Pathway-level prediction analysis

To determine which biological response programs were recovered by each model, we evaluated predictions using the 50 Hallmark gene sets from MSigDB [20]. Gene symbols were intersected with the 2,000-gene evaluation space, and pathways containing fewer than five evaluated genes were excluded.

For pathway *k*, we defined a unit-normalized membership vector *a*_*k*_ as

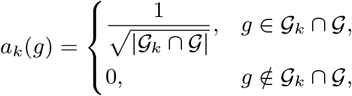

where *G* is the evaluated gene set and *G*_*k*_ is the set of genes assigned to pathway *k*.

For held-out perturbation *p*, the true and predicted centroid-relative pathway scores were defined as

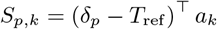

and

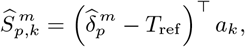

where *m* denotes the prediction model and *T*_ref_ is the reference centroid used in the corresponding evaluation setting. These scores represent the signed aggregate expression response of genes assigned to each pathway.

Pathway recovery was quantified as the Pearson correlation across perturbations:

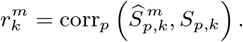

For within-cell-line analyses, out-of-fold predictions from all five cross-validation folds were concatenated before computing pathway correlations across the complete perturbation set. For cross-cell-line analyses, pathway scores were calculated from the single zero-shot target evaluation using the same target reference centroid as the perturbed-reference metric.

Model-specific pathway recovery was reported relative to the DepMap Ridge baseline as

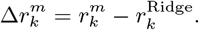

Positive values indicate stronger recovery of pathway-level response variation than DepMap Ridge, whereas negative values indicate weaker recovery.

### 4.8 Response decomposition

Response decomposition was used to separate shared response structure from perturbation-specific and context-specific variation and to determine which components were recovered by each prediction model. All decompositions were computed from observed pseudobulk responses and used exclusively for post hoc analysis. Decomposition components were not provided to models during training.

#### 4.8.1 Within-cell-line template projection

Within each cell line, perturbation responses were decomposed into a template-aligned component and a template-orthogonal residual. For each cross-validation fold, the template was estimated using only the observed responses of the training perturbations:

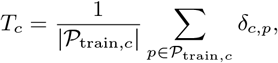

where *δ*_*c,p*_ ∈ ℝ ^*G*^ denotes the pseudobulk response to perturbation *p* in cell line *c*.

For each held-out perturbation, the true response was decomposed as

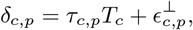

where

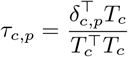

is the template coefficient and

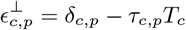

is the template-orthogonal residual.

Predicted responses were decomposed using the same observed training template:

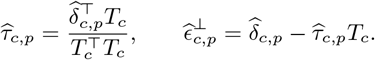

The vector *T*_*c*_ was also used as the reference centroid for perturbed-reference evaluation. However, the two operations are distinct: perturbed-reference evaluation subtracts *T*_*c*_ directly, whereas the decomposition removes only the projection of each response along the direction defined by *T*_*c*_. Estimating the template exclusively from training perturbations prevented information from held-out responses from entering the decomposition.

#### 4.8.2 Balanced cross-cell-line additive decomposition

Cross-cell-line response structure was analyzed using perturbations measured in K562, RPE1, HepG2 and Jurkat. Perturbations were represented at the target-gene level and retained only when present in all four cell lines after the uniform minimum-cell filtering described above. Responses were represented in the same ordered 2,000-gene space across datasets.

The resulting balanced response tensor contained 638 perturbations and had dimensions

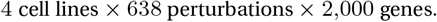

Balancing the tensor allowed cell-line, perturbation and interaction components to be estimated without confounding them with missing cell-line–perturbation combinations.

For each gene, the observed response tensor was decomposed as

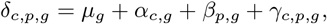

where the grand mean was

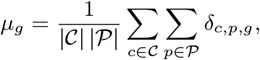

the cell-line main effect was

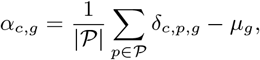

the conserved perturbation effect was

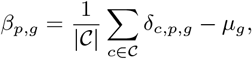

and the cell-line-by-perturbation interaction was

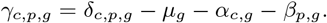

The components satisfy the zero-sum constraints

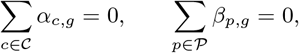

and

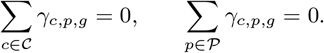

Under the balanced design, the corresponding full component tensors are mutually orthogonal under the Frobenius inner product, and their sums of squares partition the total response sum of squares.

The combined term *µ*_*g*_ + *α*_*c,g*_ is the mean perturbation response in cell line *c* and captures response structure shared across perturbations within that cell line. The conserved component *β*_*p,g*_ captures perturbation effects reproduced across cell lines, whereas *γ*_*c,p,g*_ captures cell-line-specific deviations from the conserved perturbation response.

#### 4.8.3 Template-orthogonal cross-cell-line decomposition

To characterize cross-cell-line structure after removing variation along the mean response direction, a template was calculated separately for each cell line from the balanced tensor:

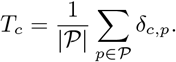

Each response was projected onto its corresponding cell-line template using

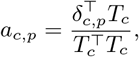

and the template-orthogonal response was defined as

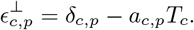

The additive decomposition was then applied to the tensor of template-orthogonal responses. Because *T*_*c*_ is the mean perturbation response within cell line *c*, the template-orthogonal tensor has zero mean across perturbations within each cell line. Its grand-mean and cell-line components therefore vanish up to numerical precision, leaving conserved perturbation and cell-line-by-perturbation interaction components.

Orthogonal projection was used for the primary analysis because it removed perturbation-to-perturbation variation in template amplitude while retaining response variation perpendicular to the template direction. Sensitivity analyses compared this operation with direct subtraction of the cell-line centroid and with ANOVA accounting, in which the *µ* + *α*_*c*_ contribution was excluded from the response-energy denominator without modifying the original responses.

#### 4.8.4 Split-half response-energy partition

Finite-cell pseudobulk responses contain sampling noise. Split-half resampling was therefore used to distinguish reproducible response structure from noise.

Within each cell-line-by-perturbation condition, cells were randomly divided into two non-overlapping subsets. Pseudobulk responses were calculated independently from the two subsets, producing response tensors Δ^(1)^ and Δ^(2)^. The complete additive decomposition was recomputed independently for each half.

Let the full component tensors be

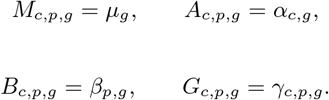

For component *Z* ∈ *{M, A, B, G}*, reproducible signal was estimated using the cross-half Frobenius inner product:

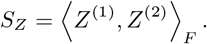

Because each component was evaluated after expansion to the full cell-line-by-perturbation-by-gene tensor, its contribution was measured on the same scale as the complete response tensor.

Under the assumption that sampling noise was independent between halves, the cross-half product retained response structure reproduced in both estimates while suppressing noise specific to either half.

The mean observed split-half response energy was

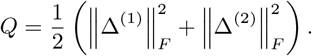

Total reproducible signal was defined as

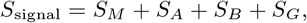

and the remaining response energy was assigned to sampling noise:

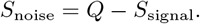

For presentation, the grand-mean and cell-line contributions were combined into a shared cell-line template term:

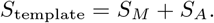

The fraction of total observed response energy assigned to each component was

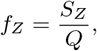

where *Z* denotes the template, conserved perturbation, interaction or noise component.

The same split-half procedure was applied to the template-orthogonal response tensor to partition residual response energy into conserved perturbation signal, cell-line-by-perturbation interaction and noise. Estimates were averaged across repeated random split-half resamples.

This procedure quantified component-level reproducibility rather than treating the complete observed interaction term as biological signal. The reproducible portion of *γ*_*c,p,g*_ represents target-specific interaction structure that is consistently detected across independent cell subsets and is therefore, in principle, available for model recovery.

### 4.9 Analysis of the decomposed spaces and biological attributions

#### 4.9.1 Template enrichment analysis

To characterize the biological programs represented by the generic perturbation template, we performed preranked gene-set enrichment analysis on the template vector estimated for each cell line.

For each cell line *c*, the template weight of gene *g* was defined as its mean response across all perturbations,

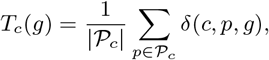

where _*c*_ denotes the set of perturbations included for cell line *c*. Genes with large positive template weights were consistently upregulated across perturbations, whereas genes with large negative template weights were consistently downregulated.

Genes were ranked by *T*_*c*_(*g*) and tested for pathway enrichment using preranked gene-set enrichment analysis. Enrichment was evaluated against the MSigDB Hallmark collection [20], comprising 50 curated gene sets representing well-defined biological states and processes. Preranked enrichment analysis was performed using `gseapy prerank` with 1,000 permutations.

The normalized enrichment score, denoted by NES, adjusts the raw enrichment score for gene-set size and the permutation-derived null distribution, allowing enrichment values to be compared across pathways and cell lines. A positive NES indicates that pathway genes were enriched among genes with positive template weights, whereas a negative NES indicates enrichment among genes with negative template weights.

False-discovery-rate-adjusted *q*-values were calculated across all tested pathways within each cell line using the Benjamini-Hochberg procedure.

#### 4.9.2 Essential versus non-essential perturbation template analysis

We used the genome-wide K562 CRISPRi Perturb-seq pseudobulk matrix described above to test whether essential-gene perturbations contributed more strongly to the generic perturbation template than non-essential perturbations. Perturbation targets were classified as essential if they appeared in the Replogle K562 essential-gene screen, yielding 1,056 essential perturbations. All remaining perturbations were classified as non-essential. Because many non-essential perturbations produced weak or undetectable transcriptional effects, the primary comparison used significant non-essential perturbations, defined by energy test *p <* 0.001, yielding 1,574 perturbations.

The essential-gene template was defined as the mean response across essential perturbations:

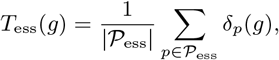

where *δ*_*p*_(*g*) is the gemgroup-normalized pseudobulk response of perturbation *p*. To compare essential and non-essential perturbations on the same response axis, all perturbations were projected onto this fixed essential-gene template. For each perturbation, the template loading was

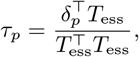

and the per-perturbation template fraction was

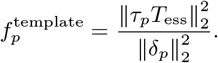

This quantity measures the fraction of each perturbation response explained by the shared essential-gene template direction. The group-level template variance fraction was computed by aggregating squared projection energy across perturbations:

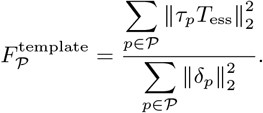

To test whether differences in template loading were explained by response magnitude, we performed response-norm matching. For each essential perturbation, the significant non-essential perturbation with the closest *L*_2_ response norm was selected without replacement, producing 1,056 matched pairs. Template loading and template fraction were then recomputed using the same essential-gene template direction. As a complementary control, essential perturbations were divided into response-norm quartiles, and significant non-essential perturbations were sampled from the same norm ranges. Template fractions were compared within each quartile.

To compare the biological identity of the essential and non-essential templates, we also computed a significant non-essential template as the mean response across significant non-essential perturbations. Genes were ranked by their signed template weights and analyzed using preranked gene-set enrichment against the MSigDB Hallmark collection [20] with 1,000 permutations. Normalized enrichment scores were compared across Hallmark pathways to determine whether the two templates represented the same biological program.

Finally, we compared response dimensionality before and after template removal. Effective rank was computed from the first 100 PCA variance components after excluding components with explained variance below 10^−12^. With normalized variance weights

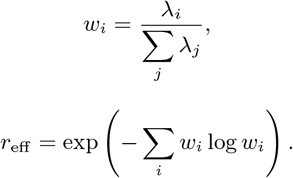

#### 4.9.3 Cross-cell-line template concordance

For the cross-cell-line analysis, template vectors were calculated independently for K562, RPE1, HepG2 and Jurkat. Preranked geneset enrichment analysis was performed separately for each cell-line template using the same Hallmark collection and enrichment settings.

The consistency of template biology between cell lines was assessed using pairwise Pearson correlations of the corresponding NES profiles. For cell lines *c* and *c*^*′*^, template concordance was calculated as

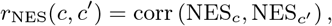

where each vector contains the enrichment scores of the 50 Hallmark pathways.

Pathways with concordant enrichment direction and magnitude across cell lines were interpreted as components of a shared perturbation-response program. Pathways showing strong enrichment in only one or a subset of cell lines were interpreted as context-dependent components of the cell-line-specific template.

#### 4.9.4 Template magnitude and variance attribution

For each cell line or perturbation group, the perturbation template was defined as the mean response vector across all perturbations in the group. For a group *P*, the template was calculated as

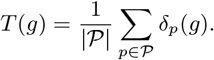

The template magnitude was quantified using its Euclidean norm,

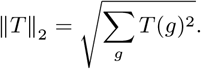

The template magnitude measures the strength of the shared perturbation response. A large value of *T* _2_ indicates that perturbations consistently shift expression in a common direction, whereas a template close to zero indicates that perturbation responses are heterogeneous and cancel when averaged.

To quantify the fraction of total perturbation response energy attributable to the template direction, the unit template vector was defined as

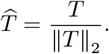

For each perturbation *p*, the template loading was calculated by projecting its response onto the unit template direction,

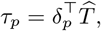

and the corresponding template-aligned response was

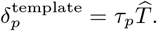

The template-removed residual was defined as

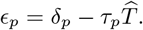

By construction, the residual was orthogonal to the template direction,

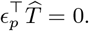

The global template fraction was defined as the ratio of template-aligned response energy to total response energy across all perturbations,

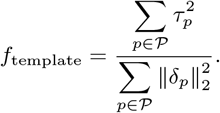

Because 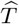 has unit norm, 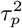 is the squared response energy of perturbation *p* along the template direction. The global template fraction therefore measures the proportion of total perturbation response energy contained in the shared template axis. Values near zero indicate that most response variation is orthogonal to the template, whereas values near one indicate that perturbations primarily differ in the magnitude of a common response pattern.

To characterize heterogeneity across individual perturbations, the per-perturbation template fraction was calculated as

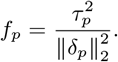

A perturbation with a high value of *f*_*p*_ produces a response dominated by the shared template direction, whereas a perturbation with a low value of *f*_*p*_ produces a response that is primarily orthogonal to the template. The distribution of *f*_*p*_ across perturbations was used to determine whether template dominance was broadly shared or concentrated among a subset of perturbations.

#### 4.9.5 Cross-cell-line template comparison

To determine whether the perturbation template represented a shared cellular response or a cell-line-specific program, templates were calculated independently for each cell line. The directional agreement between templates from cell lines *a* and *b* was quantified using cosine similarity,

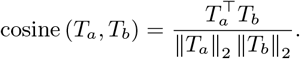

A cosine similarity close to one indicates that the same genes are consistently shifted in the same direction across the two cell lines. A value close to zero indicates that the templates occupy largely orthogonal directions, whereas a negative value indicates opposing shared response programs.

Pairwise template similarity was calculated across the four benchmark cell lines K562, RPE1, HepG2 and Jurkat. The same analysis was performed between essential and non-essential perturbation templates within the K562 genome-wide screen.

Template magnitudes were also compared across cell lines and perturbation classes to determine whether the strength of the shared response varied systematically. Within the K562 genome-wide screen, separate templates were calculated for significant essential perturbations, significant non-essential perturbations and non-significant perturbations. Significant perturbations were defined using an energy-test threshold of *p <* 0.001, whereas non-significant perturbations were defined using *p* ≥ 0.5.

The non-significant group contained perturbations whose single-cell expression distributions were not distinguishable from controls by the energy test. Its template magnitude was therefore used as an empirical reference for structured variation observed in the absence of a detectable perturbation effect.

#### 4.9.6 Control versus perturbation subspace analysis

To determine whether perturbation response variation occupied the same transcriptomic subspace as control cell-state variation, we performed reciprocal subspace projection analysis. Non-significant perturbations, defined by an energy-test value of *p* ≥ 0.5, were used as a proxy for control variation. These responses capture structured expression differences arising from biological heterogeneity, including cell-cycle and metabolic state, together with technical variation such as batch and gem-group effects, without a detectable perturbation response.

Let *X*_ctrl_ denote the matrix of non-significant perturbation responses, with perturbations represented as rows and genes as columns. Singular value decomposition was applied as

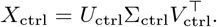

The first *k* right singular vectors defined the control subspace,

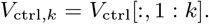

For a perturbation response matrix *X*_pert_, containing either essential or significant non-essential perturbations, the projection onto the control subspace was calculated as

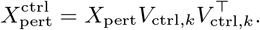

The fraction of perturbation response energy explained by the control subspace was then calculated as

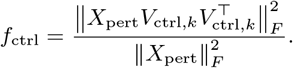

The complementary fraction,

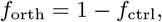

quantifies perturbation response energy orthogonal to the control subspace.

The reciprocal analysis was performed by constructing a perturbation subspace from the leading right singular vectors of *X*_pert_ and projecting the non-significant response matrix onto this subspace. This analysis tested whether control variation occupied the same axes as perturbation responses, rather than only whether perturbation responses could be represented using control-derived axes.

To distinguish structured perturbation-specific variation from measurement noise, the component orthogonal to the control subspace was calculated as

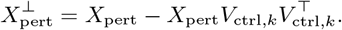

Singular value decomposition was then applied to the orthogonal residual,

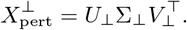

The singular values of the residual matrix were compared with a Marchenko–Pastur noise threshold for pseudobulk responses. The threshold was calculated as

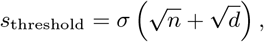

where *n* is the number of perturbations, *d* is the number of genes and *σ* is the estimated per-entry noise standard deviation. To account for the reduction in sampling variance produced by pseudobulk averaging, *σ* was approximated as

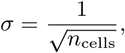

where *n*_cells_ is the number of cells contributing to each pseudobulk response.

Singular values exceeding *s*_threshold_ were interpreted as structured perturbation-specific variation beyond both the control subspace and the expected pseudobulk sampling noise. This analysis produced a three-part decomposition of perturbation response variation into variation shared with the control subspace, structured variation specific to perturbation responses and unstructured measurement noise.

#### 4.9.7 Response subspace overlap

Subspace overlap between two perturbation-response groups was quantified using the leading right singular vectors of their response matrices. Let *V*_*A,k*_ and *V*_*B,k*_ denote matrices containing the first *k* right singular vectors for response groups *A* and *B*, respectively. Their normalized subspace overlap was calculated as

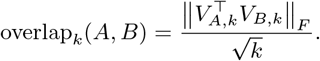

This metric is equivalent to the root mean square of the cosines of the principal angles between the two subspaces. It equals one when the two *k*-dimensional subspaces are identical and approaches zero when they are orthogonal.

Subspace overlap was calculated between essential and significant non-essential perturbation responses, between essential and non-significant responses, and between significant non-essential and non-significant responses. The analysis was performed using *k* = 10, *k* = 20 and *k* = 50 dimensions.

Comparison across these perturbation classes determined whether significant essential and non-essential perturbations activated similar transcriptomic programs and whether either response space overlapped with the structured variation observed among non-significant, control-like perturbations.

### 4.10 Prediction attribution

The fixed response decompositions were subsequently used as diagnostic bases for determining which response components were recovered by model predictions. All models were evaluated using pseudobulk response vectors in the same 2,000-gene feature space. The ground-truth decomposition components were never used as model inputs or optimization targets.

#### 4.10.1 Within-cell-line template and residual attribution

Within-cell-line attribution separately measured whether a model recovered variation in the amplitude of the shared response template and whether it recovered perturbation-specific deviations from that template.

Template coefficient recovery was calculated as the Pearson correlation between the true and predicted projection coefficients across held-out perturbations,

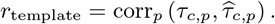

Residual recovery was calculated for each held-out perturbation as the Pearson correlation across genes between its true and predicted template-removed responses,

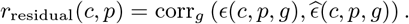

Residual correlations were then averaged across held-out perturbations,

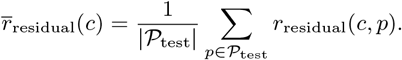

We additionally quantified the fraction of predicted response energy assigned to the template and residual subspaces. The predicted template fraction was

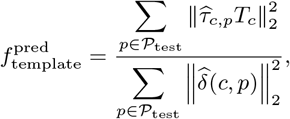

and the predicted residual fraction was

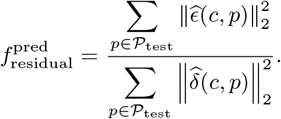

Because the template projection and residual were orthogonal,

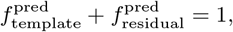

up to numerical precision.

Together, these quantities distinguished models that primarily reproduced the shared perturbation template from models that also recovered perturbation-specific residual structure.

Within-cell-line attribution was performed under the same matched five-fold perturbation-level cross-validation used for the main prediction comparison. Ridge, MLP and MORPH were evaluated on identical held-out perturbations and, where required, on the subset with available perturbation descriptors.

#### 4.10.2 Cross-cell-line conserved and interaction attribution

Cross-cell-line attribution used the conserved perturbation and interaction components obtained from the balanced ground-truth tensor. For a held-out target cell line *c* and perturbation *p*, conserved-component alignment was defined as

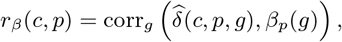

and target-interaction alignment was defined as

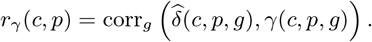

Correlations were calculated across genes for each perturbation and then averaged across perturbations in the evaluation set,

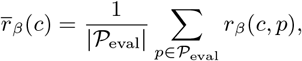

and

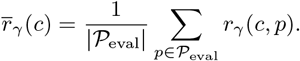

Positive alignment with *β*_*p*_(*g*) indicates that the predicted response contains perturbation structure conserved across cell lines. This is the component that can, in principle, be learned from responses measured in source cell lines.

Positive alignment with *γ*(*c, p, g*) indicates recovery of the target-specific deviation from the conserved perturbation response. These two measures therefore distinguish transfer of shared perturbation effects from prediction of how those effects are modified in the target cellular context.

The *β*_*p*_(*g*) and *γ*(*c, p, g*) components used for attribution were calculated exclusively from ground-truth responses in the balanced tensor. They were used only after prediction to interpret model outputs and were not accessible during model fitting.

#### 4.10.3 Attribution across prediction models and calibration settings

The same response construction and attribution procedure was applied to every model. Ridge and MLP predictions were obtained from the DepMap-based response-mapping models described above. Within-cell-line MORPH predictions were obtained from matched five-fold perturbation-level cross-validation.

For cross-cell-line analyses, MORPH predictions were evaluated in the one-source-to-one-target transfer setting, including source-only transfer, fine-tuning on target control cells and calibration using 30% of target perturbations. STATE predictions were evaluated in the three-source-to-one-target setting, including zero-shot transfer and calibration using 30% of target perturbations.

When target perturbations were used for calibration, attribution and prediction metrics were calculated on the same remaining 70% of target perturbations.

All single-cell model outputs were aggregated into pseudobulk response vectors using the same response construction pipeline before attribution. Differences in template, residual, conserved-component or interaction recovery therefore reflected differences in model predictions rather than differences in preprocessing or evaluation.

#### 4.10.4 Target perturbations attribution

For target calibration analyses, we compared three training regimes: zero-shot cross-cell-line prediction using source cell lines only, pooled training on source cell lines plus 30% target perturbations and target-only training using only the 30% target calibration set. In all cases, the same remaining 70% of target perturbations were used for evaluation.

For MORPH, we compared source-only transfer, target-control fine-tuning and fine-tuning with 30% target perturbations. Source-only transfer applied the source-trained model directly to target control cells. Target-control fine-tuning used reconstruction loss on target non-targeting cells with linearly annealed KL divergence. Fine-tuning with 30% target perturbations used the same target-control reconstruction objective together with supervised prediction loss on 30% of target perturbations, and evaluated performance on the remaining 70%.

For STATE, we compared source-only cross-cell-line transfer with training that additionally included 30% target-cell-line perturbations. STATE used one-hot perturbation encoding over the union perturbation vocabulary and was evaluated on the held-out target perturbations.

We applied the *β*_*p*_(*g*) and *γ*(*c, p, g*) attribution metrics to each calibration setting. This determined whether target perturbation measurements improved prediction by increasing recovery of the conserved perturbation component or by revealing the target-specific interaction. When multiple seeds were available, attribution results were averaged across seeds; Ridge was deterministic.

### 4.11 Statistical analysis and reporting

All model comparisons were performed on matched evaluation units whenever possible. Within-cell-line comparisons between Ridge, MLP and MORPH used the same perturbation-level folds, the same held-out perturbations and the same perturbed-reference evaluation metric. Cross-cell-line comparisons used matched target-cell-line perturbation sets within each transfer setting. For calibration analyses, all models used the same target calibration and test splits.

For within-cell-line five-fold cross-validation, performance was computed separately for each fold and then averaged across folds within each cell line. When multiple random seeds were used, performance was first averaged across seeds for each fold or evaluation setting, and then summarized across folds or target cell lines as appropriate. Ridge models were deterministic for a fixed split and therefore were not seed-averaged.

Pairwise model comparisons in the matched within-cell-line setting were performed using paired two-sided *t*-tests over matched folds within each cell line. We also performed pooled paired comparisons across the 20 cell-line-by-fold observations when comparing models across all four cell lines. These tests were used as descriptive comparisons of matched evaluation folds rather than as independent biological replicates.

For cross-cell-line transfer, results were reported separately by directed transfer pair, target cell line and training regime. In three-source-to-one-target experiments, each of the four cell lines was held out as the target in turn. In one-source-to-one-target experiments, results were reported for each directed source-target pair. For double-unseen experiments, performance was summarized separately for perturbations that remained observed in source cell lines and perturbations held out from all source cell lines.

For target cell line calibration analyses, results were reported for the same held-out 70% of target perturbations across zero-shot, source-plus-target-calibration and target-only training regimes. This allowed changes in performance and component attribution to be interpreted as effects of adding target perturbation measurements rather than changes in the evaluation set.

For combinatorial perturbation prediction on the Norman benchmark, performance was summarized separately for the 0*/*2, 1*/*2 and 2*/*2 unseen-gene strata. Additive-composition baselines were reported only in the 0*/*2 unseen stratum, where both constituent single-gene perturbations were observed during training and the baseline was defined. Ridge, MLP and MORPH were evaluated on the same DepMap-covered test double perturbations whenever the corresponding model was defined.

For pathway-level prediction, pathway recovery was computed independently for each retained Hallmark gene set as the Pearson correlation between true and predicted pathway activity across held-out perturbations. Model differences were reported relative to the DepMap Ridge baseline. Pathway-level statistics were used as descriptive summaries of biological response recovery and were not interpreted as independent tests across unrelated biological hypotheses.

Unless otherwise stated, reported correlations are Pearson perturbation-specific correlations computed across genes for each perturbation and then averaged across perturbations. For pairwise similarity analyses, associations between similarity matrices were quantified using Spearman correlation over upper-triangular matrix entries. For response decomposition and variance attribution analyses, reported fractions were computed from sums of squares in the corresponding response space. Split-half estimates were averaged across 50 random splits.

## Code availability

Code required to reproduce the analyses can be found at: github.com/xinyizhanglab/perturbation-decomposition.

## Author contributions

A.M. designed the research, developed and implemented the computational framework, performed model and data analysis, and wrote the paper. X.Z. designed and supervised the research, and wrote the paper.

## Acknowledgements

A.M. and X.Z. were supported by AITHYRA funding.

## Competing interests

The authors declare no competing interests.

## A. Supplementary Material

### A.1 Supplementary Figures

### A.2 Supplementary Note 1. Sensitivity of cross-cell-line response decomposition to template removal

For the balanced four-cell-line analysis, we decomposed each response into a global mean response, a cell-line component, a conserved perturbation component and a cell-line-by-perturbation interaction:

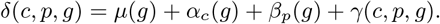

The balanced response matrix contained perturbations and genes shared across K562, RPE1, HepG2 and Jurkat. Because pseudobulk responses are estimated from finite numbers of cells, we estimated reproducible signal using split-half resampling. In each resample, cells from each cell-line-by-perturbation condition were split into two halves, pseudobulk responses were recomputed independently, and the decomposition was applied to each half. Reproducible signal for each component was estimated using the cross-half dot product between matching components. Measurement noise was defined as the remaining variance not explained by reproducible signal.

In the full response space, reproducible variance was distributed across the global and cell-line template, the conserved perturbation component and the interaction component. The global mean and cell-line component together explained 27.9% of total response variance. The conserved perturbation component explained 29.4%, the cell-line-by-perturbation interaction explained 23.5%, and the remaining 19.2% was attributable to measurement noise (Supplementary Table 1). Thus, full response structure contained both broad cell-line response programs and substantial perturbation-specific signal conserved across contexts.

We then asked whether the residual decomposition depended on how the shared template was removed. We compared three related approaches. First, in ANOVA accounting, we decomposed the full response tensor and defined the residual denominator by excluding the global mean and cell-line component. Second, in centroid removal, we subtracted the mean response of each cell line before applying the same additive decomposition. These two approaches gave nearly identical residual fractions, with approximately 40.7% to 40.8% of residual variance explained by the conserved perturbation component, 32.6% by the interaction component and 26.6% to 26.7% by measurement noise.

Third, we used projective template removal, which matches the within-cell-line template decomposition used in the main analysis. For each cell line, each perturbation response was projected onto the cell-line template direction and this projection was subtracted before applying the cross-cell-line decomposition. This stricter procedure removed all response variation along the template direction. After projective template removal, the conserved perturbation component explained 35.0% of residual variance, the cell-line-by-perturbation interaction explained 27.7%, and measurement noise explained 37.4%.

Across all three approaches, template-independent responses contained both a substantial conserved perturbation component and a substantial context-specific interaction. Projective removal reduced the total residual signal and increased the relative contribution of measurement noise, as expected from a stricter removal of the shared template direction. However, it did not eliminate either the conserved perturbation component or the interaction component. These analyses support the conclusion that residual perturbation responses contain both transferable perturbation signal and target context-dependent signal.

**Table 1.** Sensitivity of residual decomposition to template removal strategy.

| Response space / method | Template | Conserved perturbation $\beta$ | Interaction $\gamma$ | Noise |
| --- | --- | --- | --- | --- |
| Full response ANOVA | 27.9% | 29.4% | 23.5% | 19.2% |
| Residual by ANOVA accounting | — | 40.7% | 32.6% | 26.7% |
| Residual after centroid removal | — | 40.8% | 32.6% | 26.6% |
| Residual after projective template removal | — | 35.0% | 27.7% | 37.4% |

### A.3 Supplementary Note 2. Template and residual geometry of perturbation responses

We decomposed each within-cell-line pseudobulk perturbation response into a shared template component and a perturbation-specific residual (Methods). For each perturbation *p*, the response was defined as the difference between the mean expression of perturbed cells and the mean expression of non-targeting control cells from the same cell line. The template *T* was defined as the training perturbed centroid, and each response was projected onto this direction. The remaining orthogonal component was treated as the perturbation-specific residual. This decomposition allowed us to compare the geometry of the full response space with the geometry remaining after removing the shared response direction.

The contribution of the template varied substantially across cell lines (Fig. 1d, Supplementary Table 2). In K562, the template captured approximately 23% of full response energy and had norm 1.04. In RPE1, the template captured approximately 65% of full response energy and had norm 3.70. In HepG2, the template captured approximately 49% of full response energy and had norm 2.35. In Jurkat, the template captured approximately 28% of full response energy and had norm 1.48. Thus, RPE1 and HepG2 were the most template-dominated datasets, whereas K562 and Jurkat contained broader response structure outside the shared direction.

Template dominance compressed the apparent dimensionality of the full response space (Supplementary Fig. 1a, Supplementary Table 2). In RPE1, the first full response PC explained 53.3% of variance, and one PC was sufficient to explain 50% of full delta variance. In HepG2, the first response PC explained 43.1% of variance, and two PCs were sufficient to explain 50% of full delta variance. In K562 and Jurkat, the first response PC explained 21.3% and 26.7% of variance, respectively, and four PCs were required to explain 50% of full delta variance. The Shannon effective rank of the full response space was 10.8 in RPE1, 14.1 in HepG2, 25.6 in K562 and 24.1 in Jurkat.

After template removal, these differences became much smaller. The Shannon effective rank of the residual space was 32.3 in K562, 37.1 in RPE1, 31.1 in HepG2 and 34.4 in Jurkat. The first residual PC explained 20.7%, 17.1%, 20.9% and 16.0% of residual variance in K562, RPE1, HepG2 and Jurkat, respectively (Supplementary Table 2-3). The number of PCs required to explain 50% of residual variance was six in K562, seven in RPE1, five in HepG2 and six in Jurkat. These results show that apparent differences in response complexity across cell lines are largely driven by the strength of the template in the full response space. Once the template is removed, the perturbation-specific response has comparable rank across datasets.

**Table 2.** Template strength across cell lines.

| Cell line | Template variance explained | Template norm |
| --- | --- | --- |
| K562 | 23% | 1.04 |
| RPE1 | 65% | 3.70 |
| HepG2 | 49% | 2.35 |
| Jurkat | 28% | 1.48 |

**Table 3.** Full and template-removed response dimensionality.

| Cell line | Full delta effective rank | Full delta PC1 variance | PCs for 50% full delta | Residual effective rank | Residual PC1 variance | PCs for 50% residual |
| --- | --- | --- | --- | --- | --- | --- |
| K562 | 25.6 | 21.3% | 4 | 32.3 | 20.7% | 6 |
| RPE1 | 10.8 | 53.3% | 1 | 37.1 | 17.1% | 7 |
| HepG2 | 14.1 | 43.1% | 2 | 31.1 | 20.9% | 5 |
| Jurkat | 24.1 | 26.7% | 4 | 34.4 | 16.0% | 6 |

We characterized the biological content of the template using curated signed gene signatures (Methods, Supplementary Table 4). The strongest single aligned signature among those tested was proliferation shutdown, with cosine similarities of 0.270 in K562, 0.477 in RPE1, 0.441 in HepG2 and 0.279 in Jurkat. A combined p53 arrest signature showed a similar pattern, with cosine similarities of 0.219 in K562, 0.382 in RPE1, 0.378 in HepG2 and 0.194 in Jurkat. These values indicate that the template contains a reproducible growth-arrest component, strongest in RPE1 and HepG2, but is not fully explained by any single curated signature. Additional stress and cell-line-specific programs contribute to the average response.

**Table 4.**
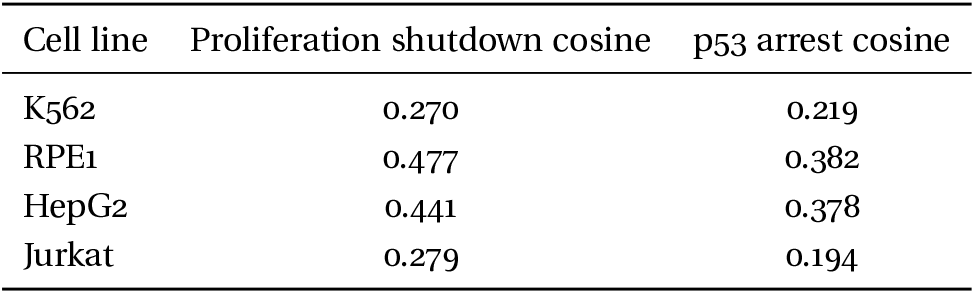
Alignment between template and curated response signatures.

We also compared the template with the principal axes of each cell-line response space (Methods). This analysis was used to measure whether the template occupied the dominant variance directions of the perturbation response space, rather than to assign a unique biological interpretation to each principal component. This distinction is important because lower PCs are constrained to be orthogonal to earlier components and can mix several biological programs. The template was strongly aligned with the leading full response PC in all cell lines, with cosines of 0.96 in K562, 0.99 in RPE1, 0.98 in HepG2 and 0.94 in Jurkat. However, the fraction of full response variance explained by this leading axis differed across cell lines. RPE1 and HepG2 had strong template-dominated leading axes, whereas K562 and Jurkat distributed response variance across more components. Thus, standard losses are expected to emphasize the template most strongly when the template is both large and concentrated in the leading response direction.

The template was not equivalent to cell identity (Supplementary Fig. 1). Cell-type-specific signatures were largely orthogonal to the average perturbation response. The K562 template showed little alignment with erythroid differentiation, even though erythroid genes contributed to the leading response structure of K562. The HepG2 template showed little alignment with hepatocyte identity. Epithelial and mesenchymal signatures were also weak across cell lines. These analyses indicate that the template is better interpreted as a perturbation-induced growth-arrest and stress response than as a lineage-marker program.

### A.4 Supplementary Note 3. Projection of perturbation responses onto control expression subspaces

To compare perturbation response geometry with natural cell state variation, we fitted PCA on centered non-targeting control cells from each cell line and projected the template and residual responses onto the resulting control subspace (Methods). This analysis asked which parts of the perturbation response were already represented by homeostatic variation.

The template was strongly represented in control space (Supplementary Fig. 1, Supplementary Table 5). Using the first 100 control PCs, the control subspace captured 92.0% of the K562 template, 98.3% of the RPE1 template, 96.8% of the HepG2 template and 95.2% of the Jurkat template. In RPE1 and HepG2, the template was already concentrated in the first control PC, which captured 74.2% and 69.8% of the template, respectively. In K562 and Jurkat, the first control PC captured only 15.8% and 11.5% of the template, respectively, but the first five control PCs captured 72.3% and 79.1%. Thus, the template is broadly contained in control variation, although its embedding in control space differs by cell line.

The residual response showed the opposite behavior. Using the first 100 control PCs, only 36.9% of K562 residual variance, 34.5% of RPE1 residual variance, 39.3% of HepG2 residual variance and 38.6% of Jurkat residual variance was captured. Conversely, 59.4% to 65.4% of residual variance remained outside the control subspace. This was consistent across all four cell lines, indicating that most template-independent response variance is not represented by the leading axes of natural control cell variation.

**Table 5.** Alignment between template and response or control principal axes.

| Cell line | $\cos(\text{template, full delta PC1})$ | $\cos(\text{template, control PC1})$ |
| --- | --- | --- |
| K562 | 0.96 | 0.40 |
| RPE1 | 0.99 | 0.86 |
| HepG2 | 0.98 | 0.84 |
| Jurkat | 0.94 | 0.34 |

**Table 6.** Template energy captured by control PCs.

| Cell line | Control PC1 | First 5 control PCs | First 100 control PCs |
| --- | --- | --- | --- |
| K562 | 15.8% | 72.3% | 92.0% |
| RPE1 | 74.2% | 88.8% | 98.3% |
| HepG2 | 69.8% | 86.3% | 96.8% |
| Jurkat | 11.5% | 79.1% | 95.2% |

We further decomposed the residual into the component inside the control subspace and the component outside the control subspace (Methods). The component inside the control subspace was comparatively compact. Across cell lines, this component accounted for approximately 35% to 42% of residual variance and required only about three PCs to explain 50% of its variance. The component outside the control subspace accounted for approximately 58% to 65% of residual variance and was substantially higher rank, requiring approximately 46 to 51 PCs to explain 50% of its variance. This suggests that the small residual fraction overlapping control variation consists of a few shared state-like modes, whereas the larger outside component contains many perturbation-induced programs.

We then compared control PCs, perturbation PCs and random bases using the subspace energy overlap metric (Methods, Supplementary Table 6-10). On full deltas, control PCs captured substantial response variance because they recovered much of the shared template. At five dimensions, control PCs captured 28.1% of full delta variance in K562, 59.1% in RPE1, 47.3% in HepG2 and 31.3% in Jurkat. The corresponding perturbation PCs captured 47.9%, 74.7%, 64.5% and 48.3% of full delta variance. Random bases captured less than 0.5% in all cases.

After template removal, the difference between control and perturbation residual subspaces became much larger. At five dimensions, control PCs captured 13.4% of residual variance in K562, 7.1% in RPE1, 11.2% in HepG2 and 11.9% in Jurkat. In contrast, residual PCs captured 34.4%, 32.1%, 34.9% and 31.2% of residual variance in the same cell lines. Random bases captured approximately 0.3%. Thus, control variation and perturbation-specific responses are not unrelated random directions, but they organize expression variation into different subspaces. Control variation captures the template well, while the residual response requires perturbation-induced directions that are poorly represented by the leading control manifold.

### A.5 Supplementary Note 4. Raw DepMap similarity is not response aligned

We asked whether DepMap coessentiality profiles define a response aligned neighborhood structure. For each cell line, we compared pairwise cosine similarity between DepMap profiles with pairwise similarity between perturbation responses. We performed this analysis both on full pseudobulk deltas and on template removed residuals (Methods, Supplementary Tables 11-13).

Raw DepMap similarity was partially correlated with full delta similarity, but this relationship disappeared after template removal. Spearman correlations between DepMap cosine similarity and full response similarity were 0.226 in K562, 0.362 in RPE1, 0.366 in HepG2 and 0.210 in Jurkat. After template removal, the corresponding correlations were close to zero or slightly negative: − 0.038 in K562, − 0.021 in RPE1, − 0.023 in HepG2 and − 0.029 in Jurkat. Thus, the apparent alignment between DepMap and full transcriptional response similarity was largely explained by the shared template rather than by perturbation specific residual structure.

Binning gene pairs by DepMap similarity gave the same conclusion. Even pairs in the highest DepMap similarity decile had mean template removed response correlations close to zero. In K562, the bottom and top DepMap similarity deciles had mean residual correlations of 0.0001 and 0.0171, respectively. In RPE1, the corresponding values were 0.0021 and 0.0099. In HepG2, they were 0.0011 and 0.0152. In Jurkat, they were − 0.0006 and 0.0157. Thus, even the most coessential gene pairs were not strong transcriptional response neighbors after the shared template was removed.

We also clustered genes in DepMap space and compared within cluster and between cluster residual response correlations (Methods). Coessentiality clusters were only weakly perturbation coherent. At coarse granularity, within cluster residual correlations were 0.005 in K562, 0.009 in RPE1, 0.005 in HepG2 and 0.004 in Jurkat, close to the between cluster baseline. At finer granularity, within cluster correlations increased slightly but remained small, reaching 0.022 in K562, 0.019 in RPE1, 0.019 in HepG2 and 0.016 in Jurkat.

These results indicate that DepMap contains biological information relevant to perturbation prediction, but this information is not organized as a fine response neighbor graph or as response coherent clusters.

**Table 7.** Residual energy captured by control PCs.

| Cell line | Control PC1 | First 5 control PCs | First 100 control PCs | Residual outside 100 control PCs |
| --- | --- | --- | --- | --- |
| K562 | 1.6% | 10.9% | 36.9% | 59.4% |
| RPE1 | 0.8% | 7.4% | 34.5% | 65.4% |
| HepG2 | 0.8% | 10.5% | 39.3% | 58.4% |
| Jurkat | 4.2% | 11.1% | 38.6% | 59.3% |

**Table 8.** Rank structure of residual components inside and outside control space.

| Cell line | Residual in control variance | PCs for 50% in-control residual | Residual outside control variance | PCs for 50% outside-control residual |
| --- | --- | --- | --- | --- |
| K562 | 36.9% | 3 | 59.4% | 46 |
| RPE1 | 34.5% | 3 | 65.4% | 51 |
| HepG2 | 39.3% | 3 | 58.4% | 46 |
| Jurkat | 38.6% | 3 | 59.3% | 49 |

**Table 9.** Subspace energy overlap for full deltas.

| Cell line | Basis | $k = 1$ | $k = 5$ | $k = 10$ |
| --- | --- | --- | --- | --- |
| K562 | Control PCs | 4.7% | 28.1% | 39.5% |
| K562 | Perturbation PCs | 23.9% | 47.9% | 55.7% |
| RPE1 | Control PCs | 47.5% | 59.1% | 68.6% |
| RPE1 | Perturbation PCs | 64.5% | 74.7% | 79.2% |
| HepG2 | Control PCs | 34.1% | 47.3% | 55.0% |
| HepG2 | Perturbation PCs | 48.7% | 64.5% | 70.8% |
| Jurkat | Control PCs | 8.3% | 31.3% | 41.5% |
| Jurkat | Perturbation PCs | 27.9% | 48.3% | 55.8% |

**Table 10.** Subspace energy overlap for template-removed residuals.

| Cell line | Basis | $k = 1$ | $k = 5$ | $k = 10$ |
| --- | --- | --- | --- | --- |
| K562 | Control PCs | 1.7% | 13.4% | 24.8% |
| K562 | Residual PCs | 14.5% | 34.4% | 43.4% |
| RPE1 | Control PCs | 0.9% | 7.1% | 17.7% |
| RPE1 | Residual PCs | 12.1% | 32.1% | 43.2% |
| HepG2 | Control PCs | 0.8% | 11.2% | 20.2% |
| HepG2 | Residual PCs | 14.3% | 34.9% | 44.6% |
| Jurkat | Control PCs | 4.9% | 11.9% | 21.8% |
| Jurkat | Residual PCs | 10.2% | 31.2% | 39.9% |

**Table 11.** Pairwise DepMap similarity versus response similarity.

| Cell line | Full delta similarity | Template removed residual similarity |
| --- | --- | --- |
| K562 | 0.226 | -0.038 |
| RPE1 | 0.362 | -0.021 |
| HepG2 | 0.366 | -0.023 |
| Jurkat | 0.210 | -0.029 |

**Table 12.** Residual response similarity in low and high DepMap similarity bins.

| Cell line | Bottom DepMap similarity decile | Top DepMap similarity decile |
| --- | --- | --- |
| K562 | 0.0001 | 0.0171 |
| RPE1 | 0.0021 | 0.0099 |
| HepG2 | 0.0011 | 0.0152 |
| Jurkat | -0.0006 | 0.0157 |

**Table 13.** Residual response coherence of DepMap clusters.

| Cell line | Coarse clusters within | Coarse clusters between | Fine clusters within | Fine clusters between |
| --- | --- | --- | --- | --- |
| K562 | 0.005 | -0.001 | 0.022 | -0.001 |
| RPE1 | 0.009 | 0.001 | 0.019 | 0.002 |
| HepG2 | 0.005 | -0.001 | 0.019 | -0.000 |
| Jurkat | 0.004 | -0.001 | 0.016 | -0.001 |

#### A.6 Supplementary Note 5. Ridge and MLP maps from DepMap to perturbation response

To test whether DepMap could directly predict perturbation responses, we trained supervised models to map DepMap profiles to pseudobulk expression deltas. Each perturbation targeting gene *g* was represented by the DepMap dependency profile of that gene. DepMap profiles were PCA-reduced on the training set, and the model was trained to predict the corresponding pseudobulk perturbation delta (Methods, Supplementary Table 14). Performance was evaluated using perturbed-reference Pearson, which subtracts the training perturbed centroid from both the prediction and ground truth before computing correlation.

Because the original random split gave slightly different test coverage across models due to missing DepMap embeddings for some perturbations, we repeated the within-cell-line comparison using five-fold perturbation-level cross-validation restricted to perturbations with available DepMap embeddings. This produced matched evaluation folds for Ridge, MLP and MORPH.

A simple Ridge regression from PCA-reduced DepMap profiles to pseudobulk deltas recovered substantial template-independent signal in all four cell lines. Mean perturbed-reference correlations were 0.340 in K562, 0.494 in RPE1, 0.413 in HepG2 and 0.311 in Jurkat. A multilayer perceptron trained on the same PCA-reduced DepMap inputs achieved similar performance, reaching 0.355, 0.491, 0.416 and 0.320, respectively. MORPH also recovered positive perturbation-specific signal, reaching 0.357, 0.459, 0.399 and 0.305, but did not outperform the simpler response-aligned maps.

Ridge and MLP were not significantly different in any cell line. MORPH was significantly lower than both Ridge and MLP in RPE1, and lower than MLP in Jurkat. In K562 and HepG2, differences between MORPH and the simpler maps were not significant. Across all 20 paired cell-line-by-fold comparisons, MLP outperformed MORPH, while Ridge and MLP were not significantly different. These results indicate that the main recoverable signal is accessible through response-aligned maps from DepMap to expression space, rather than requiring a conditional generative architecture. The central requirement is the presence of a perturbation descriptor that encodes transferable functional information about the target gene.

**Table 14.** Matched five-fold comparison of DepMap-based predictors.

| Cell line | Ridge perturbed-reference Pearson | MLP perturbed-reference Pearson | MORPH perturbed-reference Pearson |
| --- | --- | --- | --- |
| K562 | 0.340 ± 0.019 | 0.355 ± 0.010 | 0.357 ± 0.012 |
| RPE1 | 0.494 ± 0.010 | 0.491 ± 0.022 | 0.459 ± 0.015 |
| HepG2 | 0.413 ± 0.013 | 0.416 ± 0.010 | 0.399 ± 0.013 |
| Jurkat | 0.311 ± 0.009 | 0.320 ± 0.005 | 0.305 ± 0.011 |
| Pooled | 0.389 ± 0.072 | 0.395 ± 0.066 | 0.380 ± 0.058 |

### A.7 Supplementary Note 6. Predictive information resides in lower variance DepMap axes

We performed a rank ablation to identify where perturbation predictive information resides within DepMap. For each rank *k*, we retained the first *k* DepMap PCs, trained Ridge regression using the same response target, and evaluated perturbed reference Pearson (Methods, Supplementary Table 15). In K562, using only the leading DepMap PC produced weak perturbation specific prediction, with perturbed reference Pearson 0.082. Adding the first five PCs increased performance to 0.257. Performance was strongest at intermediate rank, reaching 0.366 with 50 PCs and 0.377 with 200 PCs. Using the full DepMap rank decreased performance to 0.298.

**Table 15.** DepMap rank ablation in K562.

| DepMap representation | Perturbed reference Pearson |
| --- | --- |
| PC1 | 0.082 |
| First 5 PCs | 0.257 |
| First 50 PCs | 0.366 |
| First 200 PCs | 0.377 |
| Full rank | 0.298 |

These results show that perturbation predictive information is not concentrated in the dominant DepMap axis. The leading PC captures broad high variance structure in the coessentiality matrix, including tissue specificity and broad essentiality patterns, but provides little perturbation specific response prediction when used alone. In contrast, lower variance components encode finer functional distinctions between essential gene modules. Inspection of these components identified functional subsystems including mitochondrial genes, ribosome biogenesis genes, RNA processing genes and transcriptional regulators. Thus, the useful prior signal is not the dominant raw geometry of DepMap, but a set of lower variance functional axes that become predictive after response aligned projection.

### A.8 Supplementary Note 7. Learned projections rotate DepMap into response geometry

We analyzed the learned Ridge projection to determine whether it preserved raw DepMap neighborhoods or transformed them into a response aligned geometry. First, we decomposed the Ridge weight matrix using singular value decomposition. The leading predictive mode was dominated by template independent signal, with 93.6% of its response energy lying in the residual component rather than in the shared template direction. This indicates that the learned map extracted perturbation specific response structure rather than simply reconstructing the shared perturbation template.

Second, we compared pairwise similarity before and after applying the learned response map. Raw DepMap cosine similarity was poorly aligned with template removed response similarity, as shown above. After projection through the learned map, pairwise similarity became substantially more aligned with residual response similarity, reaching Spearman correlations of approximately 0.25. Thus, the learned projection does not simply preserve coessentiality neighborhoods. It rotates dependency space into perturbation response geometry, making weak functional axes more visible in the similarity structure that is relevant for transcriptional response prediction.

### A.9 Supplementary Note 8. MORPH representation stages do not improve on raw DepMap

We next examined whether MORPH learns a response aligned perturbation geometry internally. We compared the predictive content of representations along the MORPH pipeline: the raw DepMap profile, the perturbation encoder representation, the pre-attention representation, and two post-attention representations. For each representation, we fitted the same response aligned readout and evaluated perturbed reference Pearson on held out perturbations (Methods, Supplementary Table 16). This analysis asked whether MORPH progressively transforms DepMap into a representation that is more predictive of template removed transcriptional response.

The raw DepMap profile was the strongest representation in all four cell lines. It reached perturbed reference correlations of 0.343 in K562, 0.471 in RPE1, 0.392 in HepG2 and 0.308 in Jurkat. Compressing the 1,208 dimensional DepMap profile through the MORPH perturbation encoder reduced performance to 0.289, 0.419, 0.360 and 0.249, respectively. Thus, the perturbation encoder removed predictive information that was available in the raw dependency profile.

The pre-attention representation gave identical performance to the perturbation encoder representation, with perturbed reference correlations of 0.289 in K562, 0.419 in RPE1, 0.360 in HepG2 and 0.249 in Jurkat. This indicates that adding the cell embedding before attention did not add measurable perturbation response information in this representation-level analysis. The target cell context available to MORPH at this stage therefore did not make the perturbation descriptor more response aligned.

Post-attention representations also did not recover the lost signal. The first post-attention representation reached 0.299 in K562, 0.428 in RPE1, 0.350 in HepG2 and 0.246 in Jurkat. The second post-attention representation reached 0.298, 0.428, 0.348 and 0.245. These values remained below the raw DepMap profile in every cell line. Cross-attention therefore did not restore the predictive information lost during perturbation encoding and did not introduce additional response aligned structure.

Together, these analyses show that MORPH’s perturbation prediction signal is carried primarily by the DepMap descriptor itself. In this setting, the conditional VAE architecture did not improve access to that signal. Instead, the perturbation encoder compressed away part of the predictive information, the pre-attention cell representation added no measurable information, and downstream attention layers did not recover the lost response aligned signal. This supports the conclusion that the key bottleneck is not the existence of biological signal in DepMap, but how efficiently a model preserves and aligns dependency variation with template removed perturbation response structure.

**Table 16.** Predictive content of MORPH representation stages.

| Representation stage | K562 | RPE1 | HepG2 | Jurkat |
| --- | --- | --- | --- | --- |
| Raw DepMap | 0.343 | 0.471 | 0.392 | 0.308 |
| Perturbation encoder | 0.289 | 0.419 | 0.360 | 0.249 |
| Pre-attention | 0.289 | 0.419 | 0.360 | 0.249 |
| Post-attention 1 | 0.299 | 0.428 | 0.350 | 0.246 |
| Post-attention 2 | 0.298 | 0.428 | 0.348 | 0.245 |

### A.10 Supplementary Note 9. Direct source to target prediction with STATE and MORPH

We first evaluated direct transfer between K562 and RPE1 using STATE and MORPH (Methods, Supplementary Table 17). In this setting, models were trained only in the source cell line and evaluated in the target cell line. This experiment tested whether a model trained on source cell line perturbation responses could recover template removed target responses without any target perturbation measurements.

Source only transfer recovered little template removed signal in either direction. In K562 to RPE1 transfer, STATE reached 0.046 perturbed reference Pearson and MORPH reached -0.013. In RPE1 to K562 transfer, STATE reached 0.022 and MORPH reached -0.041. Standard Pearson was higher in some settings, but this reflected recovery of shared response structure rather than perturbation specific residuals.

Adding target perturbations improved both models, especially STATE. With 30% of target perturbations available during training, STATE reached 0.532 perturbed reference Pearson in K562 to RPE1 and 0.458 in RPE1 to K562. MORPH also improved, reaching 0.226 and 0.116 in the same directions. These results indicate that target perturbation measurements reveal target response structure that is not predictable from source cell line responses alone.

**Table 17.**
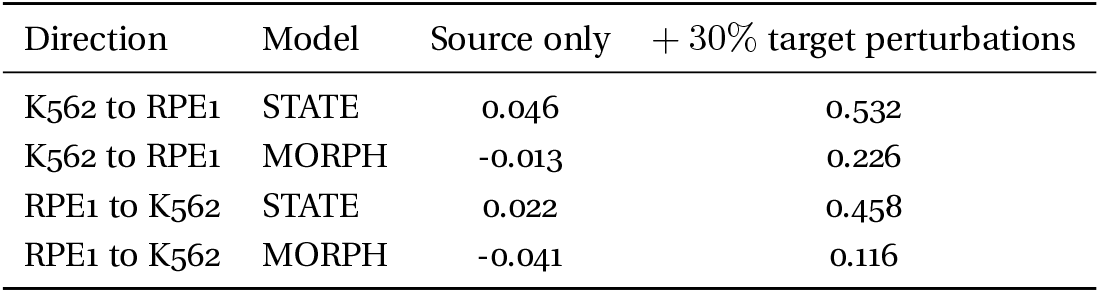
Direct K562 and RPE1 transfer with STATE and MORPH.

| Direction | Model | Source only | + 30% target perturbations |
| --- | --- | --- | --- |
| K562 to RPE1 | STATE | 0.046 | 0.532 |
| K562 to RPE1 | MORPH | -0.013 | 0.226 |
| RPE1 to K562 | STATE | 0.022 | 0.458 |
| RPE1 to K562 | MORPH | -0.041 | 0.116 |

### A.11 Supplementary Note 10. DepMap supports cross-context and double-unseen prediction

In the three-source-to-one-target zero-shot setting, MLP models recovered positive perturbation-specific signal in all four target cell lines (Supplementary Fig. 4, Supplementary Table 18). Perturbed-reference correlations were 0.226 in K562, 0.271 in RPE1, 0.340 in HepG2 and 0.241 in Jurkat. Ridge models showed the same pattern but lower performance, reaching 0.118, 0.208, 0.287 and 0.167. Thus, DepMap-based models can transfer perturbation information into a cellular context not observed during training.

In the double-unseen setting, performance decreased but remained positive across all target cell lines (Supplementary Fig. 4, Supplementary Table 18). Ridge reached perturbed-reference correlations of 0.102 in K562, 0.211 in RPE1, 0.276 in HepG2 and 0.161 in Jurkat. MLP reached 0.164, 0.231, 0.288 and 0.194, respectively. The persistence of positive signal in this setting shows that DepMap contains information about perturbations that were never observed in the training response data. The reduction relative to zero-shot context transfer shows that observing the same perturbation in source cell lines still provides useful response information.

Adding 30% target perturbations produced modest improvements in several targets. In the three-source-to-one-target setting, MLP performance increased from 0.340 to 0.386 in HepG2, from 0.226 to 0.240 in K562 and from 0.241 to 0.259 in Jurkat, while RPE1 showed no improvement. Ridge showed a similar but weaker pattern. Thus, target perturbation measurements can improve prediction, but the cross-context signal recovered from DepMap remains limited by the target response landscape.

**Table 18.** Three-source-to-one-target cross-cell-line prediction.

| Target cell line | Ridge zero shot | MLP zero shot | Ridge + 30% target | MLP + 30% target | Ridge double unseen | MLP double unseen |
| --- | --- | --- | --- | --- | --- | --- |
| K562 | 0.118 | 0.226 | 0.135 | 0.240 | 0.102 | 0.164 |
| RPE1 | 0.208 | 0.271 | 0.205 | 0.256 | 0.211 | 0.231 |
| HepG2 | 0.287 | 0.340 | 0.334 | 0.386 | 0.276 | 0.288 |
| Jurkat | 0.167 | 0.241 | 0.188 | 0.259 | 0.161 | 0.194 |

### A.12 Supplementary Note 11. One-source-to-one-target transfer reveals source-target asymmetries

We also evaluated one-source-to-one-target transfer, where only a single source cell line was available for training (Methods, Supplementary Table 19). This setting increased dependence on both the source and target contexts. In zero-shot transfer, MLP reached perturbed-reference correlations of 0.211 for K562 to RPE1, 0.128 for RPE1 to K562, 0.283 for Jurkat to HepG2 and 0.197 for HepG2 to Jurkat. Ridge followed the same ordering but was lower in most settings, reaching 0.164, 0.038, 0.235 and 0.149.

In the double-unseen setting, MLP retained positive signal for all pairs but performance decreased to 0.153 for K562 to RPE1, 0.069 for RPE1 to K562, 0.241 for Jurkat to HepG2 and 0.191 for HepG2 to Jurkat. Ridge reached 0.132, 0.024, 0.222 and 0.181. The weakest case was RPE1 to K562, consistent with K562 being a difficult target response landscape and with the limited ability of one source cell line to cover the target context.

**Table 19.** One-source-to-one-target cross-cell-line prediction.

| Source to target | Ridge zero shot | MLP zero shot | Ridge + 30% target | MLP + 30% target | Ridge double unseen | MLP double unseen |
| --- | --- | --- | --- | --- | --- | --- |
| K562 to RPE1 | 0.164 | 0.211 | 0.188 | 0.203 | 0.132 | 0.153 |
| RPE1 to K562 | 0.038 | 0.128 | 0.068 | 0.141 | 0.024 | 0.069 |
| Jurkat to HepG2 | 0.235 | 0.283 | 0.325 | 0.349 | 0.222 | 0.241 |
| HepG2 to Jurkat | 0.149 | 0.197 | 0.193 | 0.231 | 0.181 | 0.191 |

### A.13 Supplementary Note 12. Target response properties partly explain cross-context performance

Cross-cell-line performance was better explained by properties of the target cell line than by the identity of the source cell lines. In the three-source-to-one-target zero-shot setting, HepG2 was the easiest target, followed by RPE1, Jurkat and K562. This ordering was consistent with the target response geometry measured above. HepG2 and RPE1 had lower apparent response rank and stronger template structure, whereas Jurkat and K562 had broader response spaces and weaker templates (Supplementary Table 20).

This suggests that DepMap provides a transferable perturbation descriptor, but the amount of recoverable signal depends on the target response landscape. A cell line with a lower rank or more template-dominated response space is easier to predict into because a larger fraction of its response can be explained by shared or conserved components. A cell line with broader residual geometry requires more target-specific information, reducing the amount of response signal that can be inferred from DepMap alone.

**Table 20.**
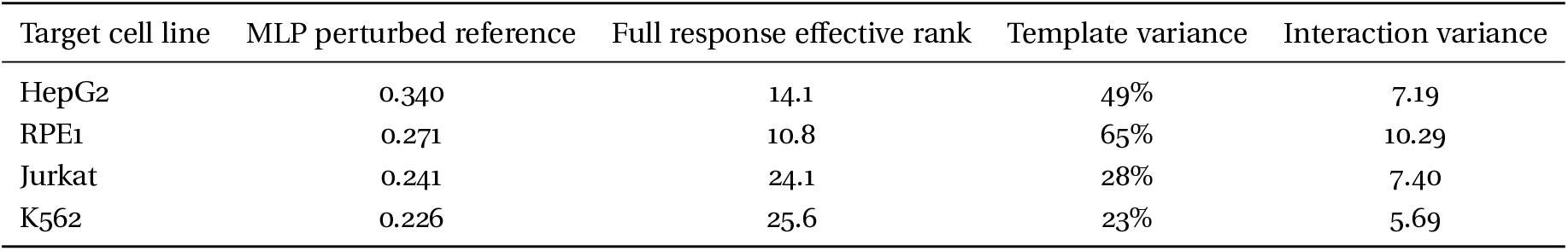
Target properties and three-source zero-shot MLP performance.

| Target cell line | MLP perturbed reference | Full response effective rank | Template variance | Interaction variance |
| --- | --- | --- | --- | --- |
| HepG2 | 0.340 | 14.1 | 49% | 7.19 |
| RPE1 | 0.271 | 10.8 | 65% | 10.29 |
| Jurkat | 0.241 | 24.1 | 28% | 7.40 |
| K562 | 0.226 | 25.6 | 23% | 5.69 |

### A.14 Supplementary Note 13. Response-basis gaps quantify the barrier for program-transfer architectures

To measure the difficulty of transferring response programs across cell lines, we computed how much target residual variance was spanned by the source cell line response basis (Methods, Supplementary Table 21). At rank 20, source response programs captured only 20% to 30% of target residual variance across cell line pairs. The corresponding basis gap was 0.766 for K562 to RPE1, 0.779 for RPE1 to K562, 0.802 for Jurkat to HepG2, 0.758 for HepG2 to Jurkat, 0.711 for K562 to Jurkat and 0.702 for HepG2 to RPE1.

These large gaps are consistent with the weak cross cell line overlap of residual perturbation spaces reported above. They show that architectures that transfer perturbation effects through a shared source response basis face a substantial context barrier. Most target residual variance is not spanned by the source response programs, even when the same perturbations and genes are available in both cell lines.

However, response-basis gap did not explain the performance of DepMap based gene level operators. Jurkat to HepG2 had the largest basis gap but was among the best one-to-one zero shot transfers, whereas RPE1 to K562 had a similar basis gap but was the weakest transfer. This shows that response-program transfer and DepMap-to-response prediction are distinct mechanisms. Program-transfer architectures are limited by source-target residual subspace overlap. DepMap based operators instead learn a direct mapping from perturbation function to gene level response, making performance depend more on target response complexity and on the predictable component of the target interaction term.

**Table 21.** Response-basis gaps and one-to-one zero-shot prediction.

| Source to target | Basis gap | Basis overlap | MLP perturbed reference | Ridge perturbed reference |
| --- | --- | --- | --- | --- |
| K562 to RPE1 | 0.766 | 23.4% | 0.211 | 0.164 |
| RPE1 to K562 | 0.779 | 22.1% | 0.128 | 0.038 |
| Jurkat to HepG2 | 0.802 | 19.8% | 0.283 | 0.235 |
| HepG2 to Jurkat | 0.758 | 24.2% | 0.197 | 0.149 |
| K562 to Jurkat | 0.711 | 28.9% | — | — |
| HepG2 to RPE1 | 0.702 | 29.8% | — | — |

### A.15 Supplementary Note 14. Reference baselines for target-only and within-cell-line prediction

For reference, we compared cross-cell-line prediction with target-only calibration and full within-cell-line prediction (Methods, Supplementary Table 22). In the target-only 30% setting, the model was trained only on 30% of target perturbations and evaluated on the remaining target perturbations, with no source cell line data. MLP reached perturbed-reference correlations of 0.307 in K562, 0.425 in RPE1, 0.402 in HepG2 and 0.262 in Jurkat. Ridge reached 0.306, 0.466, 0.397 and 0.276.

Full within-cell-line training provided the strongest reference for most cell lines. MLP reached 0.356 in K562, 0.458 in RPE1, 0.372 in HepG2 and 0.318 in Jurkat, while Ridge reached 0.343, 0.469, 0.393 and 0.309. These reference results show that target perturbation measurements remain the most direct source of target response information. DepMap supports cross-context and double-unseen extrapolation, but it does not fully replace target response observations.

**Table 22.** Reference baselines.

| Setting | Cell line | Ridge | MLP |
| --- | --- | --- | --- |
| Target-only 30% | K562 | 0.306 | 0.307 |
| Target-only 30% | RPE1 | 0.466 | 0.425 |
| Target-only 30% | HepG2 | 0.397 | 0.402 |
| Target-only 30% | Jurkat | 0.276 | 0.262 |
| Within cell line | K562 | 0.343 | 0.356 |
| Within cell line | RPE1 | 0.469 | 0.458 |
| Within cell line | HepG2 | 0.393 | 0.372 |
| Within cell line | Jurkat | 0.309 | 0.318 |

### A.16 Supplementary Note 15. Target calibration saturates after approximately 100 to 150 perturbations

We quantified how many target perturbations were needed to calibrate target response geometry (Methods). Ridge based coordinate mapping improved rapidly as target perturbations were added, increasing from near zero with one calibration perturbation to perturbed reference Pearson values of approximately 0.38 to 0.40 after 100 to 150 target perturbations (Supplementary Fig. 4). Beyond this range, additional target perturbations produced diminishing returns, with less than 0.01 improvement per 50 added perturbations.

This saturation is consistent with a low dimensional response space view. If the dominant target response geometry can be represented by a limited number of response programs, then a moderate number of target perturbations is sufficient to estimate much of the response program alignment and gene level readout. Target calibration therefore does not require observing all perturbations, but it does require enough interventions to identify how perturbation programs are expressed in the target cellular context.

### A.17 Supplementary Note 16. Prediction decomposition reveals which response components models learn

To determine which response components were recovered by each model, we applied the same response decomposition used in the main text to model predictions. Within each cell line, predicted responses were decomposed into a shared template coefficient and a template removed residual. For a predicted response 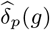, we projected onto the training perturbation centroid *T* (*g*) to obtain the predicted template coefficient 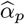, and defined the predicted residual 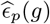 as the component orthogonal to this template direction. We then compared 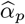 with the true template coefficient *α*_*p*_, and 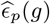 with the true template removed residual *ϵ*_*p*_(*g*). We also measured the fraction of predicted variance assigned to the template direction and to the residual component.

Within cell lines, DepMap based predictors recovered the shared template more strongly than the perturbation specific residual (Supplementary Table 23). Ridge models showed strong template coefficient recovery, with correlations between predicted and true template coefficients ranging from 0.63 to 0.75. Residual recovery was weaker but consistently positive, with residual correlations between 0.21 and 0.26. Prediction variance was also strongly template dominated, especially in RPE1 and HepG2, where 94.0% and 82.2% of Ridge predicted variance lay in the template direction. MLP predictions shifted more variance toward the residual in K562 and RPE1, but retained substantial template structure. These results show that positive perturbation prediction performance can arise from a mixture of strong template recovery and weaker perturbation specific residual recovery.

**Table 23.** Within cell line prediction decomposition.

| Cell line | Model | Perturbed reference Pearson | $\text{corr}(\hat{\alpha}, \alpha)$ | $\text{corr}(\hat{\epsilon}, \epsilon)$ | Template variance | Residual variance |
| --- | --- | --- | --- | --- | --- | --- |
| K562 | Ridge | 0.345 | 0.626 | 0.257 | 55% | 45% |
| K562 | MLP | 0.350 | 0.615 | 0.249 | 45% | 55% |
| RPE1 | Ridge | 0.470 | 0.750 | 0.209 | 94% | 6% |
| RPE1 | MLP | 0.458 | 0.737 | 0.219 | 86% | 14% |
| HepG2 | Ridge | 0.393 | 0.708 | 0.215 | 82% | 18% |
| Jurkat | Ridge | 0.309 | 0.672 | 0.214 | 72% | 28% |

We next asked whether these components could be recovered from baseline cellular context features rather than from DepMap. We trained the same prediction models using six control expression descriptors, using DepMap alone, or using DepMap concatenated with control expression descriptors (Methods, Supplementary Table 24). Control expression descriptors alone recovered little perturbation specific signal. Perturbed reference correlations were near zero in all four cell lines, and predicted variance was almost entirely template dominated. Adding control expression descriptors to DepMap did not improve performance and did not change the template residual attribution. Thus, the perturbation specific signal recovered by DepMap based models was not explained by simple baseline expression descriptors.

**Table 24.** Within cell line feature ablation and prediction attribution.

| Cell line | Feature | Perturbed reference Pearson | $\text{corr}(\hat{\alpha}, \alpha)$ | $\text{corr}(\hat{\epsilon}, \epsilon)$ | Template variance | Residual variance |
| --- | --- | --- | --- | --- | --- | --- |
| K562 | DepMap PCA50 | 0.344 | 0.628 | 0.255 | 55.4% | 44.6% |
| K562 | Control expression | -0.031 | 0.003 | -0.015 | 97.8% | 2.2% |
| K562 | DepMap + control expression | 0.343 | 0.628 | 0.254 | 55.3% | 44.7% |
| RPE1 | DepMap PCA50 | 0.470 | 0.752 | 0.207 | 94.0% | 6.0% |
| RPE1 | Control expression | 0.015 | 0.121 | 0.013 | 99.3% | 0.7% |
| RPE1 | DepMap + control expression | 0.469 | 0.750 | 0.205 | 94.0% | 6.0% |
| HepG2 | DepMap PCA50 | 0.391 | 0.703 | 0.214 | 82.2% | 17.8% |
| HepG2 | Control expression | 0.081 | 0.092 | -0.003 | 99.3% | 0.7% |
| HepG2 | DepMap + control expression | 0.390 | 0.705 | 0.214 | 82.4% | 17.6% |
| Jurkat | DepMap PCA50 | 0.310 | 0.675 | 0.216 | 71.8% | 28.2% |
| Jurkat | Control expression | 0.026 | -0.102 | 0.013 | 97.2% | 2.8% |
| Jurkat | DepMap + control expression | 0.309 | 0.670 | 0.214 | 71.8% | 28.2% |

For cross cell line prediction, we decomposed model predictions according to the balanced cell line by perturbation decomposition described in the main text and Methods. Using the perturbations shared across the four cell lines, we compared each prediction with the conserved perturbation component *β*_*p*_(*g*) and the target specific interaction *γ*(*c, p, g*). This analysis separated the response component that is transferable across cell lines from the component that depends on how a perturbation is expressed in the target cell line.

In the three source to one target zero shot setting, DepMap based predictors recovered part of the conserved perturbation component but not the target specific interaction (Supplementary Table 25). MLP predictions showed stronger recovery of *β*_*p*_(*g*) than Ridge predictions, with *R*^2^(*β*) between 0.25 and 0.39 compared with 0.10 to 0.13 for Ridge. However, correlations with *γ*(*c, p, g*) were negative across all targets. Thus, the MLP advantage in cross cell line prediction reflected better recovery of conserved perturbation signal, not recovery of the target specific interaction.

**Table 25.**
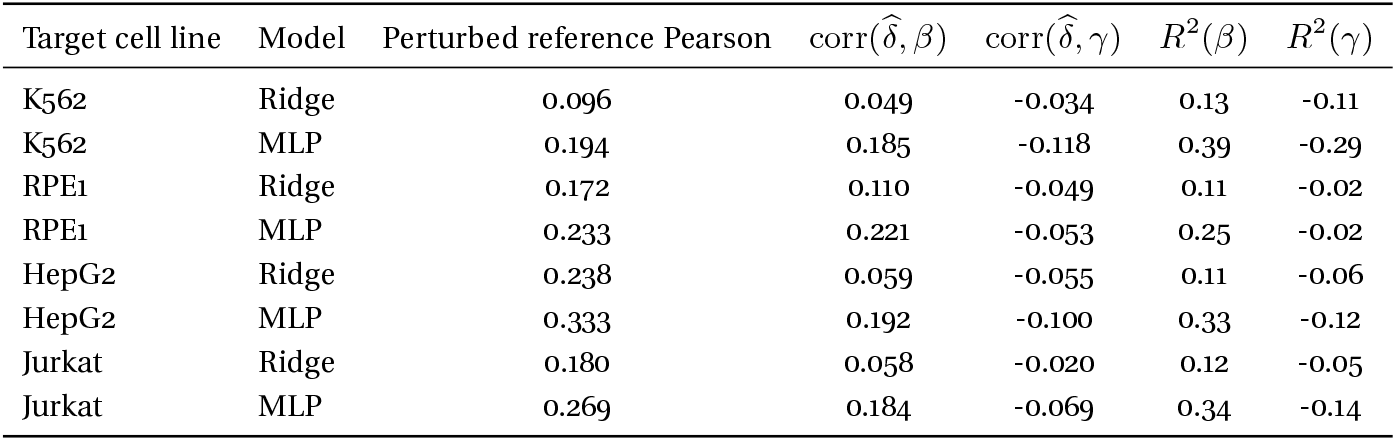
Cross cell line prediction decomposition for Ridge and MLP.

We also tested whether control expression descriptors from the target or source cell lines could provide the missing target context information. They did not. In the cross cell line setting, target control descriptors alone produced weak predictions and did not recover the conserved perturbation component. Concatenating target or source control descriptors with DepMap left perturbed reference performance and component alignment essentially unchanged (Supplementary Table 26). Therefore, the target context information required for cross cell line prediction is not captured by these baseline expression descriptors. The missing information is the target perturbation response geometry, namely the mapping from perturbation function to cell line specific transcriptional response.

**Table 26.** Cross cell line feature ablation.

| Target cell line | Feature | Perturbed reference Pearson | $\text{corr}(\hat{\delta}, \beta)$ | $\text{corr}(\hat{\delta}, \gamma)$ |
| --- | --- | --- | --- | --- |
| K562 | DepMap PCA50 | 0.096 | 0.050 | -0.035 |
| K562 | Control expression from target cell line | -0.048 | -0.128 | 0.057 |
| K562 | DepMap + target control expression | 0.096 | 0.048 | -0.034 |
| K562 | DepMap + source control expression | 0.095 | 0.048 | -0.033 |
| RPE1 | DepMap PCA50 | 0.172 | 0.110 | -0.049 |
| RPE1 | Control expression from target cell line | 0.083 | -0.126 | -0.044 |
| RPE1 | DepMap + target control expression | 0.172 | 0.110 | -0.047 |
| RPE1 | DepMap + source control expression | 0.171 | 0.109 | -0.049 |
| HepG2 | DepMap PCA50 | 0.238 | 0.060 | -0.055 |
| HepG2 | Control expression from target cell line | 0.037 | -0.126 | -0.011 |
| HepG2 | DepMap + target control expression | 0.238 | 0.060 | -0.055 |
| HepG2 | DepMap + source control expression | 0.239 | 0.059 | -0.054 |
| Jurkat | DepMap PCA50 | 0.181 | 0.058 | -0.020 |
| Jurkat | Control expression from target cell line | -0.005 | -0.125 | 0.034 |
| Jurkat | DepMap + target control expression | 0.180 | 0.057 | -0.020 |
| Jurkat | DepMap + source control expression | 0.180 | 0.055 | -0.019 |

We then examined how target calibration changed the recovered response components (Supplementary Table 27). In zero shot prediction, correlations with the interaction term were negative across targets. Training on 30% target perturbations without source data flipped the interaction correlation from negative to positive in K562, HepG2 and Jurkat, while the conserved perturbation component changed little or decreased. Pooling source and target data produced little improvement relative to zero shot. These results indicate that target perturbation measurements improve prediction by revealing the target cell line by perturbation interaction, rather than by improving recovery of the conserved perturbation component. They also show that source responses can dominate simple pooled models and obscure the weaker target calibration signal.

We applied the same prediction decomposition to STATE and MORPH (Supplementary Table 28-29. For STATE, we evaluated a three source to one target setting with no target perturbation data, and the same setting after adding 30% target perturbations. In the zero shot setting, STATE recovered the template coefficient but almost no perturbation specific residual or target interaction. Perturbed reference correlations were close to zero across targets, residual correlations were near zero, and correlations with *γ*(*c, p, g*) were weak or negative. Adding 30% target perturbations improved performance and shifted the interaction component toward positive alignment, most clearly in RPE1 and HepG2. Thus, target perturbation data revealed target response behavior that was not predictable from source cell lines alone.

For MORPH, we evaluated one source to one target transfer pairs, target control fine tuning, and fine tuning with 30% target perturbations. Source trained MORPH models retained some template alignment but recovered little perturbation specific residual, little conserved perturbation signal and no positive target interaction. Fine tuning on target controls alone did not change this behavior. In contrast, fine tuning with 30% target perturbations strongly improved perturbed reference performance, residual recovery, conserved perturbation alignment and target interaction alignment. Thus, MORPH benefits from target perturbation measurements because they reveal how the DepMap perturbation descriptor should be converted into a target cell line response, whereas target baseline information alone is insufficient.

**Table 27.** Effect of target calibration on conserved and interaction components.

| Target cell line | Setting | $\text{corr}(\hat{\delta}, \beta)$ | $\text{corr}(\hat{\delta}, \gamma)$ | Perturbed reference Pearson |
| --- | --- | --- | --- | --- |
| K562 | Zero shot | 0.050 | -0.031 | 0.103 |
| K562 | Source + 30% target pooled | 0.052 | -0.028 | 0.109 |
| K562 | Target only 30% | 0.065 | 0.039 | 0.245 |
| HepG2 | Zero shot | 0.060 | -0.048 | 0.242 |
| HepG2 | Source + 30% target pooled | 0.057 | -0.041 | 0.253 |
| HepG2 | Target only 30% | 0.034 | 0.053 | 0.323 |
| Jurkat | Zero shot | 0.058 | -0.023 | 0.175 |
| Jurkat | Source + 30% target pooled | 0.056 | -0.018 | 0.183 |
| Jurkat | Target only 30% | 0.023 | 0.051 | 0.264 |

**Table 28.** STATE prediction decomposition with and without target perturbation calibration.

| Target cell line | Setting | Perturbed reference Pearson | $\text{corr}(\hat{\alpha}, \alpha)$ | $\text{corr}(\hat{\epsilon}, \epsilon)$ | $\text{corr}(\hat{\delta}, \beta)$ | $\text{corr}(\hat{\delta}, \gamma)$ |
| --- | --- | --- | --- | --- | --- | --- |
| K562 | Zero shot | 0.034 | 0.663 | 0.009 | 0.043 | -0.021 |
| RPE1 | Zero shot | 0.058 | 0.648 | -0.017 | 0.120 | 0.014 |
| HepG2 | Zero shot | 0.077 | 0.724 | -0.003 | 0.082 | -0.013 |
| Jurkat | Zero shot | 0.045 | 0.633 | 0.028 | 0.059 | -0.021 |
| K562 | +30% target perturbations | 0.105 | 0.791 | 0.057 | 0.093 | 0.010 |
| RPE1 | +30% target perturbations | 0.367 | 0.922 | 0.081 | 0.210 | 0.259 |
| HepG2 | +30% target perturbations | 0.216 | 0.870 | 0.080 | 0.186 | 0.106 |
| Jurkat | +30% target perturbations | 0.086 | 0.628 | 0.048 | 0.070 | 0.016 |

**Table 29.** MORPH prediction decomposition across transfer and fine tuning settings.

| Transfer pair | Setting | Perturbed reference Pearson | $\text{corr}(\hat{\alpha}, \alpha)$ | $\text{corr}(\hat{\epsilon}, \epsilon)$ | $\text{corr}(\hat{\delta}, \beta)$ | $\text{corr}(\hat{\delta}, \gamma)$ |
| --- | --- | --- | --- | --- | --- | --- |
| K562 to RPE1 | Source only | 0.022 | 0.233 | -0.008 | 0.030 | -0.023 |
| RPE1 to K562 | Source only | 0.002 | 0.436 | 0.018 | 0.047 | -0.051 |
| HepG2 to Jurkat | Source only | 0.008 | 0.516 | 0.016 | 0.013 | -0.015 |
| Jurkat to HepG2 | Source only | 0.044 | 0.453 | 0.011 | 0.063 | -0.020 |
| K562 to RPE1 | Target controls only | 0.031 | 0.275 | -0.011 | 0.031 | -0.015 |
| RPE1 to K562 | Target controls only | -0.005 | 0.438 | 0.020 | 0.028 | -0.040 |
| HepG2 to Jurkat | Target controls only | 0.000 | 0.414 | 0.021 | -0.005 | -0.018 |
| Jurkat to HepG2 | Target controls only | 0.047 | 0.282 | 0.012 | 0.060 | -0.020 |
| K562 to RPE1 | +30% target perturbations | 0.412 | 0.694 | 0.275 | 0.278 | 0.206 |
| RPE1 to K562 | +30% target perturbations | 0.369 | 0.537 | 0.330 | 0.292 | 0.051 |
| HepG2 to Jurkat | +30% target perturbations | 0.396 | 0.707 | 0.322 | 0.251 | 0.110 |
| Jurkat to HepG2 | +30% target perturbations | 0.412 | 0.737 | 0.334 | 0.268 | 0.119 |

Together, these supplementary analyses support the prediction decomposition described in the main text. Within cell lines, models recover the template more strongly than the perturbation specific residual. Across cell lines, models recover part of the conserved perturbation component but fail to recover the target specific interaction in zero shot settings. Control expression descriptors and target control fine tuning do not provide this missing interaction information. Target perturbation measurements help because they reveal the cell line by perturbation interaction that is not predictable from source cell lines or baseline expression features alone.

## Notes

https://github.com/xinyizhanglab/perturbation-decomposition

### Competing Interest Statement

The authors have declared no competing interest.

## References

[1] Dixit, A., Parnas, O., Li, B., Chen, J., Fulco, C. P., Jerby-Arnon, L., Marjanovic, N. D., Dionne, D., Burks, T., Raychowdhury, R., et al. “Perturb-Seq: dissecting molecular circuits with scalable single-cell RNA profiling of pooled genetic screens”. In: cell 167.7 (2016), pp. 1853–1866 (cit. on p. 1).

[2] Adamson, B., Norman, T. M., Jost, M., Cho, M. Y., Nuñez, J. K., Chen, Y., Villalta, J. E., Gilbert, L. A., Horlbeck, M. A., Hein, M. Y., et al. “A multiplexed single-cell CRISPR screening platform enables systematic dissection of the unfolded protein response”. In: Cell 167.7 (2016), pp. 1867–1882 (cit. on p. 1).

[3] Datlinger, P., Rendeiro, A. F., Schmidl, C., Krausgruber, T., Traxler, P., Klughammer, J., Schuster, L. C., Kuchler, A., Alpar, D., and Bock, C. “Pooled CRISPR screening with single-cell transcriptome readout”. In: Nature methods 14.3 (2017), pp. 297–301 (cit. on p. 1).

[4] Replogle, J. M., Saunders, R. A., Pogson, A. N., Hussmann, J. A., Lenail, A., Guna, A., Mascibroda, L., Wagner, E. J., Adelman, K., Lithwick-Yanai, G., et al. “Mapping information-rich genotype-phenotype landscapes with genome-scale Perturb-seq”. In: Cell 185.14 (2022), pp. 2559–2575 (cit. on pp. 1, 2, 10).

[5] Nadig, A., Replogle, J. M., Pogson, A. N., Murthy, M., McCarroll, S. A., Weissman, J. S., Robinson, E. B., and O’Connor, L. J. “Transcriptome-wide analysis of differential expression in perturbation atlases”. In: Nature Genetics 57.5 (2025), pp. 1228–1237 (cit. on pp. 1, 2, 10).

[6] Roohani, Y., Huang, K., and Leskovec, J. Predicting transcriptional outcomes of novel multigene perturbations with GEARS. 2024 (cit. on pp. 1, 13).

[7] Lotfollahi, M., Klimovskaia Susmelj, A., De Donno, C., Hetzel, L., Ji, Y., Ibarra, I. L., Srivatsan, S. R., Naghipourfar, M., Daza, R. M., Martin, B., et al. “Predicting cellular responses to complex perturbations in high-throughput screens”. In: Molecular systems biology 19.6 (2023), MSB202211517 (cit. on p. 1).

[8] Hetzel, L., Boehm, S., Kilbertus, N., Günnemann, S., Theis, F., et al. “Predicting cellular responses to novel drug perturbations at a single-cell resolution”. In: Advances in Neural Information Processing Systems 35 (2022), pp. 26711–26722 (cit. on p. 1).

[9] Cui, H., Wang, C., Maan, H., Pang, K., Luo, F., Duan, N., and Wang, B. “scGPT: toward building a foundation model for single-cell multiomics using generative AI”. In: Nature methods 21.8 (2024), pp. 1470–1480 (cit. on pp. 1, 2, 9, 13).

[10] Adduri, A. K., Gautam, D., Bevilacqua, B., Imran, A., Shah, R., Naghipourfar, M., Teyssier, N., Ilango, R., Nagaraj, S., Dong, M., et al. “Predicting cellular responses to perturbation across diverse contexts with State”. In: BioRxiv (2025), pp. 2025–06 (cit. on pp. 1, 3, 4, 9, 14).

[11] He, C., Zhang, J., Dahleh, M., and Uhler, C. “MORPH Predicts the Single-Cell Outcome of Genetic Perturbations Across Conditions and Data Modalities”. In: bioRxiv (2025) (cit. on pp. 1–4, 9, 14).

[12] Ahlmann-Eltze, C., Huber, W., and Anders, S. “Deep-learning-based gene perturbation effect prediction does not yet outperform simple linear baselines”. In: Nature Methods 22.8 (2025), pp. 1657–1661 (cit. on pp. 1, 2, 9).

[13] Viñas Torné, R., Wiatrak, M., Piran, Z., Fan, S., Jiang, L., Teichmann, S. A., Nitzan, M., and Brbić, M. “Systema: a framework for evaluating genetic perturbation response prediction beyond systematic variation”. In: Nature Biotechnology (2025), pp. 1–10 (cit. on pp. 1, 4, 9, 12).

[14] Tsherniak, A., Vazquez, F., Montgomery, P. G., Weir, B. A., Kryukov, G., Cowley, G. S., Gill, S., Harrington, W. F., Pantel, S., Krill-Burger, J. M., et al. “Defining a cancer dependency map”. In: Cell 170.3 (2017), pp. 564–576 (cit. on pp. 1, 9, 10, 12).

[15] Replogle, J. M., Norman, T. M., Xu, A., Hussmann, J. A., Chen, J., Cogan, J. Z., Meer, E. J., Terry, J. M., Riordan, D. P., Srinivas, N., et al. “Combinatorial single-cell CRISPR screens by direct guide RNA capture and targeted sequencing”. In: Nature biotechnology 38.8 (2020), pp. 954–961 (cit. on pp. 3, 10, 16).

[16] Gaudelet, T., Del Vecchio, A., Carrami, E. M., Cudini, J., Kapourani, C.-A., Uhler, C., and Edwards, L. “Season combinatorial intervention predictions with Salt & Peper”. In: arXiv preprint arXiv:2404.16907 (2024) (cit. on pp. 3, 16).

[17] Zhang, J., Ubas, A. A., De Borja, R., Svensson, V., Thomas, N., Thakar, N., Lai, I., Winters, A., Khan, U., Jones, M. G., et al. “Tahoe-100m: A giga-scale single-cell perturbation atlas for context-dependent gene function and cellular modeling”. In: BioRxiv (2025), pp. 2025–02 (cit. on pp. 3, 9, 10, 17).

[18] Rogers, D. and Hahn, M. “Extended-connectivity fingerprints”. In: Journal of chemical information and modeling 50.5 (2010), pp. 742–754 (cit. on pp. 3, 9, 11, 17).

[19] Duran-Frigola, M., Pauls, E., Guitart-Pla, O., Bertoni, M., Alcalde, V., Amat, D., Juan-Blanco, T., and Aloy, P. “Extending the small-molecule similarity principle to all levels of biology with the Chemical Checker”. In: Nature Biotechnology 38.9 (2020), pp. 1087–1096 (cit. on pp. 3, 9, 11, 17).

[20] Liberzon, A., Birger, C., Thorvaldsdóttir, H., Ghandi, M., Mesirov, J. P., and Tamayo, P. “The molecular signatures database hallmark gene set collection”. In: Cell systems 1.6 (2015), pp. 417–425 (cit. on pp. 11, 19, 22, 23).

[21] Glorot, X., Bordes, A., and Bengio, Y. “Deep sparse rectifier neural networks”. In: Proceedings of the fourteenth international conference on artificial intelligence and statistics. JMLR Workshop and Conference Proceedings. 2011, pp. 315–323 (cit. on p. 13).

[22] Kingma, D. P. and Ba, J. “Adam: A Method for Stochastic Optimization”. In: International Conference on Learning Representations. 2015 (cit. on pp. 13, 14).

[23] Brown, T., Mann, B., Ryder, N., Subbiah, M., Kaplan, J. D., Dhariwal, P., Neelakantan, A., Shyam, P., Sastry, G., Askell, A., et al. “Language models are few-shot learners”. In: Advances in neural information processing systems 33 (2020), pp. 1877–1901 (cit. on p. 14).

